# *In Vivo* Generation of Bone Marrow from Embryonic Stem Cells in Interspecies Chimeras

**DOI:** 10.1101/2021.09.30.462528

**Authors:** Bingqiang Wen, Guolun Wang, Enhong Li, Olena A. Kolesnichenko, Zhaowei Tu, Senad Divanovic, Tanya V. Kalin, Vladimir V. Kalinichenko

**Author notes:** Correspondence to: Vladimir V. Kalinichenko, Center for Lung Regenerative Medicine, Cincinnati Children’s Hospital Medical Center, 3333 Burnet Avenue, Cincinnati, OH 45229, USA.

## Abstract

Generation of bone marrow (BM) from embryonic stem cells (ESCs) or induced pluripotent stem cells (iPSCs) promises to accelerate the development of future cell therapies for life-threatening disorders. However, such approach is limited by technical challenges to produce a mixture of functional BM progenitor cells able to replace all hematopoietic cell lineages. Herein, we used blastocyst complementation to simultaneously produce all BM hematopoietic cell lineages from mouse ESCs in a rat. Based on FACS analysis and single-cell RNA sequencing, mouse ESCs differentiated into hematopoietic progenitor cells and multiple hematopoietic cell types that were indistinguishable from normal mouse BM cells based on gene expression signatures and cell surface markers. Transplantation of ESC-derived BM cells from mouse-rat chimeras rescued lethally-irradiated syngeneic mice and resulted in long-term contribution of donor cells to hematopoietic cell lineages. Altogether, a fully functional bone marrow was generated from mouse ESCs using rat embryos as “bioreactors”.

**KEY POINTS:** - We used blastocyst complementation to simultaneously produce all bone marrow hematopoietic cell lineages from mouse ESCs in a rat.

- ESC-derived cells from mouse-rat chimeras were fully functional and exhibited normal gene expression signatures and cell surface markers.

## INTRODUCTION

The bone marrow (BM) is a remarkably complex organ consisting of multiple mesenchymal, immune, endothelial, and neuronal cell types which together comprise a highly specialized microenvironment required to support life-long blood regeneration or hematopoiesis (1–5). Hematopoiesis occurs in a stepwise manner and is initiated by a heterogeneous, multipotent, population of hematopoietic stem cells (HSCs), located at the apex of the hematopoietic differentiation tree. Long-term HSCs (LT-HSCs) remain quiescent to maintain their undifferentiated state within the bone marrow niche. When necessary, LT-HSCs can either undergo differentiation or self-renewal, to maintain the HSC pool. Conversely, short-term HSCs (ST-HSCs) are restricted in their self-renewal capacity and primed for differentiation into multipotent progenitors (MPPs), initiating the process of blood cell development. MPPs further differentiate into common myeloid progenitors (CMPs), lymphoid-primed multipotent progenitors (LMPPs) and common lymphoid progenitors (CLPs) that become increasingly lineage restricted with subsequent cell divisions, ultimately yielding all mature blood cell types (6). The complexities of the hematopoietic system have been studied extensively *in vitro*, utilizing paired-daughter and colony forming unit (CFU) assays (4, 5). Fluorescence-activated cell sorting (FACS) has allowed for precise isolation and characterization of HSCs and progenitor populations based on cell surface markers. Classically, the most biologically relevant way to test HSC function remains to be through serial transplantation and hematopoietic reconstitution of irradiated recipient mice (4, 5, 7). Recent advances in single cell RNA sequencing (scRNAseq) have made it possible to further explore heterogeneity of the bone marrow niche (2, 3), and identify gene expression signatures of hematopoietic progenitor cells as they differentiate into mature blood cell types (1, 8).

Generation of functional bone marrow from embryonic stem cells (ESCs) or induced pluripotent stem cells (iPSCs) will provide new therapeutic opportunities for hematologic and autoimmune disorders. However, this approach is limited by technical challenges to produce functional HSCs or the mixture of hematopoietic progenitors capable of replacing all mature blood cell types after cell transplantation. HSC-like cells have been generated from mouse and human ESCs and iPSCs using *in vitro* differentiation protocols (9–15). Likewise, ESCs and iPSCs have been used to produce myeloid and lymphoid progenitor cells as well as differentiated hematopoietic cells, including neutrophils, monocytes, erythroid cells, T and B lymphocytes (15–21). When transplanted into irradiated animals, ESC/iPSC-derived hematopoietic progenitor cells undergo differentiation and engraft into the bone marrow niche, providing an important source of renewal and regeneration for various blood cell lineages (4, 5, 14). While ESC/iPSC-derived hematopoietic cells often express appropriate cell markers, gene expression and functional studies indicate significant differences between ESC/iPSC-derived cells and endogenous cells that have undergone normal morphogenesis in the bone marrow (14, 22, 23).

*In vivo* differentiation of ESCs into multiple cell lineages can be achieved using blastocyst complementation, in which donor ESCs are injected into blastocysts of recipient animals to create chimeras. Fluorescently labeled ESCs undergo differentiation in recipient embryos that serve as “biological reactors” by providing growth factors, hormones and cellular niches to support ESC differentiation in the embryo. In mouse and rat apancreatic *Pdx1^-/-^* embryos, donor ESCs formed an entire pancreas in which both exocrine and endocrine cells were almost entirely derived from ESCs or iPSCs (24, 25). Mouse ESC/iPSC-derived β-cells from mouse-rat chimeras were fully differentiated and successfully rescued syngeneic diabetic mice (25). ESCs generated pancreatic cell lineages in apancreatic pigs (26), kidney in *Sall1*- deficient rats (27), endothelial cells in *Flk1*^-/-^ mice (28), lymphocytes in immunodeficient mice (29) and neuronal progenitors in mice with forebrain-specific overexpression of diphtheria toxin (30). Recently, mouse ESCs were used to generate lung and thyroid tissues in embryos deficient for *Fgf10*, *Nkx2-1*, *Fgfr2* or *β-catenin* (31–33). Using blastocyst complementation, mouse ESCs effectively produced hematopoietic cells in mice deficient for *Kit* or *Flk1* (28, 34). ESC-derived endothelial progenitor cells from mouse-rat chimeras were indistinguishable from endogenous endothelial progenitor cells based on gene expression signatures and functional properties (35). While all these studies support the effectiveness of blastocyst complementation for differentiation of multiple cell types from ESCs/iPSCs *in vivo*, generation of functional bone marrow from ESCs in interspecies chimeras has not yet been achieved.

Herein, we used blastocyst complementation to produce mouse bone marrow in a rat. ESC-derived cells from multiple hematopoietic cell lineages were indistinguishable from normal mouse hematopoietic BM cells based on gene expression signatures and cell surface markers. Transplantation of ESC-derived BM cells into lethally-irradiated syngeneic mice prevented mortality and resulted in a long-term contribution to BM and mature blood cell types. Our data demonstrate that interspecies chimeras can be used as “bioreactors” for *in vivo* differentiation and functional studies of ESC-derived hematopoietic progenitor cells.

## METHODS

### Ethics and data sharing statement

Bone marrow single cell RNA-seq datasets were uploaded to the Gene Expression Omnibus (GEO) database (accession number GSE184940) and made available to other investigators for purposes of replicating the procedures or reproducing the results. All animal studies were reviewed and approved by the Institutional Animal Care and Use Committee of the Cincinnati Children’s Research Foundation.

### Mice, rats and generation of mouse-rat and mouse-mouse chimeras through blastocyst complementation

C57BL/6 mice were purchased from Jackson Lab. Interspecies mouse-rat chimeras were generated using blastocyst complementation as described (35, 36). Briefly, blastocysts from SD rats were obtained at embryonic day 4.5 (E4.5), injected with fifteen GFP- labeled mouse ESC cells (ESC-GFP, C57BL/6 background) (32, 37) and transferred into pseudo pregnant SD rat females. Mouse-mouse chimeras were generated by complementing CD1 blastocysts with fifteen mouse ESC-GFP cells. For FACS analysis and bone marrow transplantation, BM cells were collected from chimeric pups that were harvested between postnatal day 4 (P4) and P10. For single cell RNA sequencing, BM cells were prepared from P10 chimeric pups. To perform BM transplantation, BM cells from 2 tibias and 2 fibulas of mouse-rat chimeras were FACS-sorted for GFP^+^ cells. 500,000 of GFP^+^ BM cells from 5-9 mouse-rat chimeras were intravenously (i.v.) injected into irradiated C57BL/6 male mice (6-8 weeks of age) via the tail vein. Whole-body irradiation (11.75 Gy) was performed 3 hours before BM transplant. Mice were harvested after 8 days or 5 months after BM transplantation. Tissue dissection, processing and preparation of single cell suspensions were carried out as described (38–42). Blood analysis was performed in animal facility of Cincinnati Children’s Hospital Research Foundation.

### Single Cell RNAseq analysis of ESC-derived bone marrow cells

Bone marrow cells from three P10 mouse-rat chimeras or three P10 mouse-mouse (control) chimeras were FACS-sorted for GFP and the *lineage* (Lin) marker. To enrich for hematopoietic progenitor cells, 90% of GFP^+^Lin^−^ cells were combined with 10% of GFP^+^Lin^+^ cells. The cell mixture was used for single cell RNA sequencing analysis based on the 10X Chromium platform (https://research.cchmc.org/pbge/lunggens/mainportal.html). RNA-seq datasets were uploaded to the GEO database (accession number GSE184940). Read alignments, quality controls and false discovery rates were described previously (43, 44). Identification of cell clusters and quantification of cluster-specific gene expression in BM scRNAseq datasets was performed as described (1, 32, 35). To assess the transcriptomic similarity of ESC-derived cells from mouse-rat and mouse-mouse BM, the scRNAseq datasets were normalized with *SCTransform* and then integrated utilizing the canonical correlation analysis (CCA). In the integrated scRNAseq datasets, the *SelectIntegrationFeatures* in Seurat package (version 4.0.0 in R 4.0 statistical environment) was used to identify anchors for integration. The *RunPCA* function was used for Principal component analysis (PCA) of scRNAseq datasets, and the *PCElbowPlot* function was used to calculate the standard deviations of the principal components (PCs). PCs with standard deviation > 3.5 were chosen as input parameters for non-linear UMAP clustering analysis. Next, the *FindNeighbors* function was used to compute the k.param nearest neighbors, and BM cell clusters were identified by a shared nearest neighbor (SNN) modularity optimization clustering algorithm implemented in the *FindClusters* function with resolution set at 0.4 (32, 35, 43).

### FACS Analysis

FACS analysis was performed using cells obtained from the bone marrow and blood. Antibodies for FACS analysis are listed in Suppl. Table S1. Immunostaining of cell suspensions were performed as described (45, 46). Identification of hematopoietic cell types based on multiple cell surface markers is described in (47–51). Stained cells were analyzed using a five-laser FACSAria II (BD Biosciences) (37, 52).

### Statistical Analysis

Statistical significance was determined using one-way ANOVA and Student’s t-test. Multiple means were compared using one-way analysis of variance with the post-hoc Tukey test. P ≤ 0.05 was considered statistically significant. For datasets with n<5, non-parametric Mann-Whitney U test was used to determine statistical significance. Data were presented as mean ± standard error of mean (SEM).

## RESULTS

### Generation of bone marrow from pluripotent embryonic stem cells in interspecies mouse-rat chimeras

To determine whether mouse ESCs can differentiate into multiple hematopoietic cell lineages in the bone marrow of a rat, blastocyst complementation was performed by injecting GFP-labeled mouse C57BL/6 ESCs (ESC-GFP) into rat SD blastocysts to create interspecies mouse-rat chimeras. Chimeric embryos were transferred into surrogate female rats for subsequent development *in utero* (Fig. 1A). While mouse-rat chimeras were viable, they were smaller than age-matched rats (Fig. 1B). Consistent with the presence of mouse ESC-derived cells (black) in the skin tissue (35), mixed black and white pigmentation distinguished the mouse-rat chimeras from juvenile rats (Fig. 1B). The average body weight of mouse-rat chimeras was smaller than rats, but larger than mice of similar age (Fig. 1C). ESC-derived cells were abundant in femur and tibia bones of the chimeras as evidenced by GFP fluorescence (Fig. 1D). FACS analysis of BM cells obtained from juvenile mouse-rat chimeras revealed that the percentage of ESC-derived cells was 15-50% (Fig. 1E-F). Thus, ESCs contribute to the bone marrow of mouse-rat chimeras.

**Figure 1.**
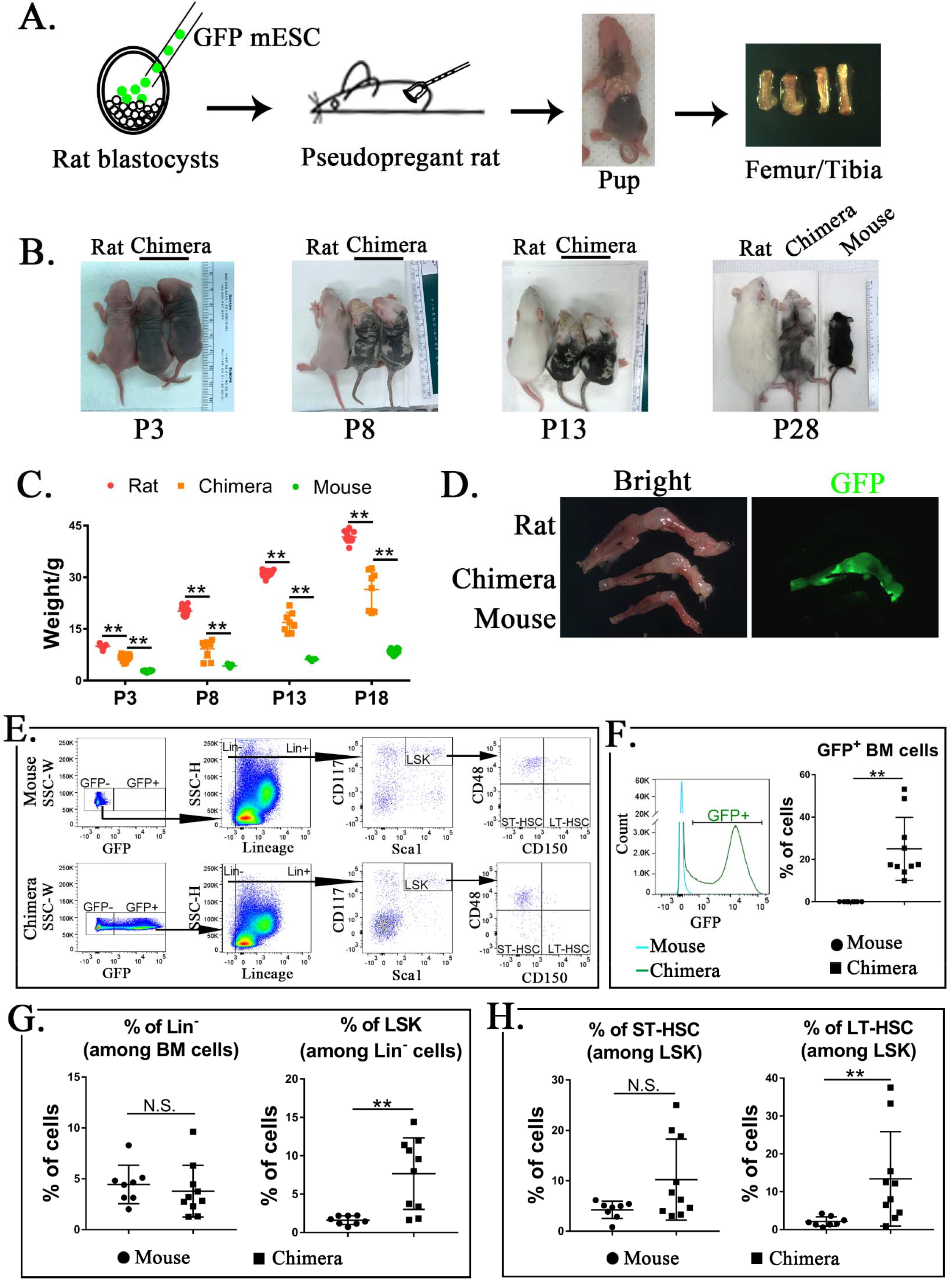
Mouse ESCs contribute to hematopoietic stem cells in the bone marrow of mouse-rat chimeras. ***A,*** Schematic shows blastocyst complementation of rat embryos with mouse ESCs to generate interspecies mouse-rat chimeras. GFP-labeled mouse ESCs (mESCs) were injected into rat blastocysts, which were implanted into surrogate rat females to undergo embryonic development in utero. Femur and tibia bones of the chimeras were used to obtain bone marrow (BM) cells. ***B,*** Photographs of mouse-rat chimeras are taken at postnatal (P) days P3, P8, P13 and P28. Mixed black and white pigmentation distinguishes the mouse-rat chimeras from juvenile rats and mice. ***C,*** Weights of mouse-rat chimeras are shown at different time points and compared to rats and mice of similar ages. Chimeras are significantly smaller than rats, but larger than mice (n=7-18 in each group), p<0.01 is **. ***D,*** Fluorescence microscopy shows GFP and bright field images of femur and tibia bones from P4 rat, mouse and mouse-rat chimera. ***E,*** FACS analysis of mouse ESC-derived (GFP-positive) cells in the bone marrow of P10 mouse-rat chimeras. Lineage-negative (Lin^−^), LSK, ST-HSC and LT-HSC cell subsets were identified in the bone marrow of mouse-rat chimeras (n=10) and control mice (n=8). ***F,*** Histograms show GFP fluorescence of BM cells from chimeras and control mice. ***G-H,*** FACS analysis show increased percentages of mouse LSKs and LT-HSCs in BM of mouse-rat chimeras (n=10) compared to control mice (n=8), p<0.01 is **, N.S. indicates no significance.

To identify ESC-derived hematopoietic stem cells (HSCs), we used GFP fluorescence and mouse-specific antibodies recognizing multiple cell surface antigens (Fig. 1E and Suppl. Fig. S1A-B). First, ESC-derived GFP^+^ BM cells were subdivided into *lineage-positive* (Lin^+^) and *lineage-negative* subpopulations (Lin^−^) (Fig. 1E and Suppl. Fig. S1A-B). The percentage of ESC-derived Lin^−^ cells in the bone marrow of mouse-rat chimeras was similar to the percentage of Lin^−^ cells in the bone marrow of age-matched C57BL/6 mice (Fig. 1E and 1G). Next, we used Sca1 and CD117 (c-KIT) antibodies to identify Lin^−^Sca1^+^c-KIT^+^ cells (LSKs) (Fig. 1E). The percentage of LSKs was higher in the bone marrow of mouse-rat chimeras compared to the control (Fig. 1G). Based on cell surface expression of CD150 and CD48, the percentage of LT-HSCs among LSKs were also higher in mouse-rat chimeras (Fig. 1E and 1H). The percentage of ST-HSCs was unchanged (Fig. 1H). Thus, mouse ESCs are capable of differentiating into hematopoietic progenitor cells in the bone marrow of mouse-rat chimeras.

### Single cell RNA sequencing identifies multiple subpopulations of ESC-derived hematopoietic cells in the bone marrow of mouse-rat chimeras

To identify ESC-derived cells in the bone marrow, single cell RNAseq (the 10X Chromium platform) of FACS-sorted GFP^+^ BM cells was performed. Mouse ESC-derived cells from P10 mouse-rat chimeras were compared to ESC-derived cells from P10 mouse-mouse (control) chimeras, the latter of which were produced by complementing mouse blastocysts with mouse ESCs from the same ESC- GFP cell line. Based on GFP fluorescence, contribution of ESCs to BM cells in both chimeras was similar (Suppl. Fig. S2A-B). To enrich for hematopoietic progenitor cells, 90% of FACS-sorted GFP^+^Lin^−^ cells were mixed with 10% of GFP^+^Lin^+^ cells prior to single cell RNA sequencing. BM cells from 3 animals per group were combined prior to FACS sorting. Based on published gene expression signatures of mouse BM cells (1), 11,326 cells from 14 major cell subtypes were identified: 5308 cells from control mouse-mouse chimeras and 6018 cells from mouse-rat chimeras (Fig. 2A). These include lymphoid, erythroid, myeloid and neutrophil progenitors, Pro-B, Pre-B, B and T lymphocytes, megakaryocytes, dendritic cells, neutrophils, basophils/eosinophils, monocytes and lymphoid-primed multipotent progenitor cells (LMPPs) (Fig. 2A and Suppl. Fig. S3A). Combined analysis of BM cells from mouse-rat and mouse-mouse chimeras demonstrated similar distributions of hematopoietic cell lineages derived from common myeloid progenitor (CMP) and common lymphoid progenitor (CLP) (Fig. 2A-B), indicating identical cell types in mouse-rat and control chimeras. For selected genes, we used violin plots to confirm cell specificity and expression levels of *Ptprc (Cd45), Pclaf, Vpreb1, Tmpo, Ebf1, Ms4a4b, Vamp5, Elof1, Elane, Ms4a2, Siglech, Ngp, Clec4d, Ctss and Ftl1-ps1* in the combined dataset (Suppl. Fig. S4). Markers of endothelial cells, adipocytes, osteocytes and neuronal cells were undetectable in BM cell suspensions from both chimeras (Suppl. Fig. S3B).

**Figure 2.**
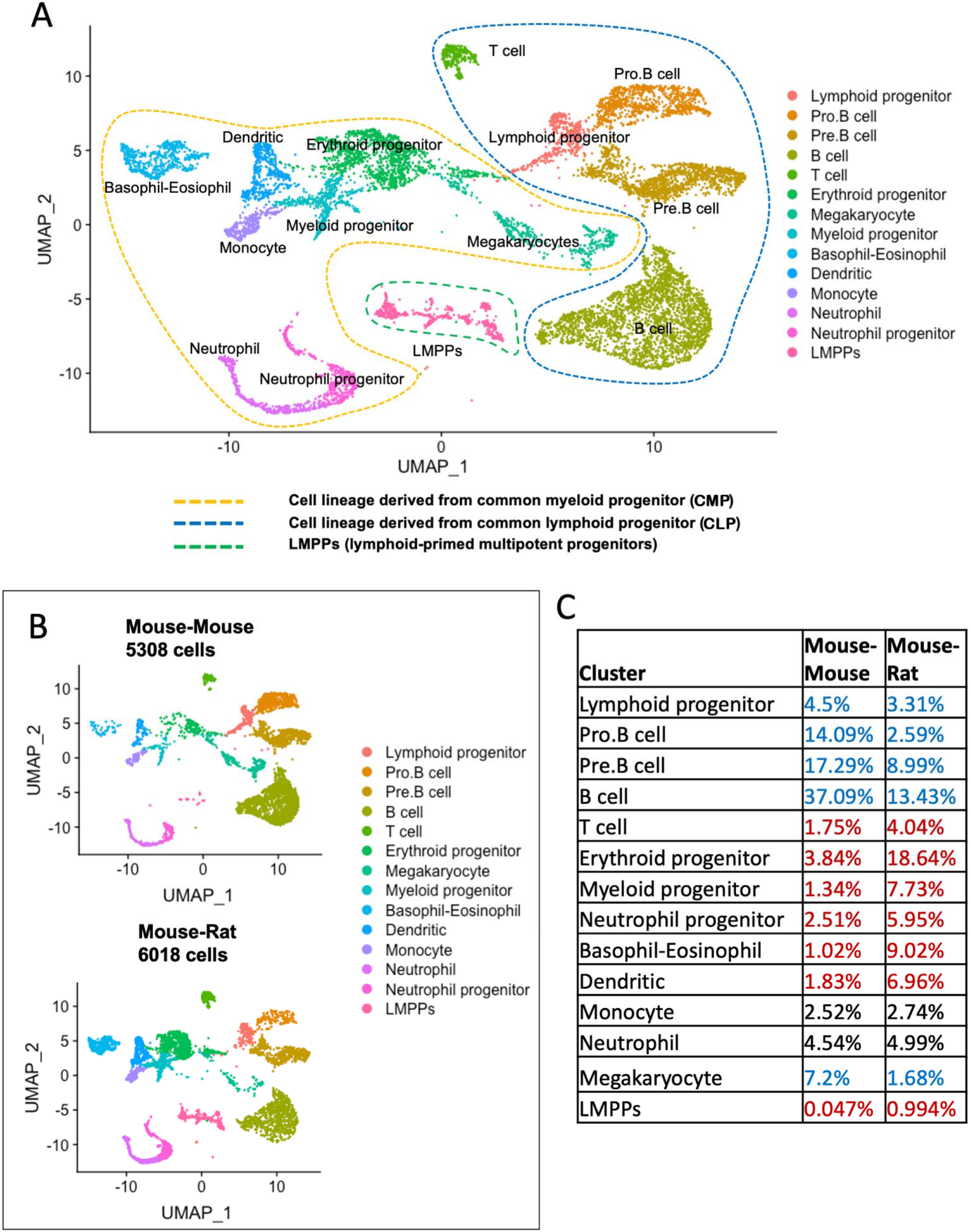
Single cell RNAseq analysis identifies ESC-derived hematopoietic cell lineages in the bone marrow of mouse-rat chimeras. ***A,*** The integrated projection of BM hematopoietic cells from mouse-rat and mouse-mouse (control) chimeras. ESC-derived BM cells were obtained from the bone marrow of P10 chimeras using FACS sorting for GFP^+^ cells. Cell clusters were identified using Uniform Manifold Approximation and Projection (UMAP) method. Cells derived from common lymphoid progenitor (CLP) are shown with blue dashed line. Cells derived from common myeloid progenitor (CMP) are shown with yellow dashed line. Cell cluster of lymphoid-primed multipotent progenitors (LMPPs) is indicated by green dashed line. ***B,*** Parallel dimension UMAP plots show identical hematopoietic cell clusters in the bone marrow of mouse-mouse chimera (5308 cells) and mouse-rat chimera (6018 cells). ***C,*** Table shows percentages of cells in individual clusters in mouse-mouse and mouse-rat chimeras. Blue color indicates decreased percentages of cells in mouse-rat chimeras compared to mouse-mouse chimeras. Red color indicates increased percentages of cells in mouse-rat chimeras.

Percentages CLP-derived lymphoid progenitors, Pro-B, Pre-B, and B cells were lower in mouse-rat chimeras compared to the control (Fig. 2C). In contrast, percentages of CMP-derived erythroid, myeloid and neutrophil progenitors, dendritic cells, and basophils/eosinophils were higher (Fig. 2C). Monocytes and neutrophils were similar, whereas megakaryocytes were decreased in the bone marrow of mouse-rat chimeras (Fig. 2C). The percentage of lymphoid-primed multipotent progenitors (LMPPs) in mouse-rat chimeras was increased compared to the control (Fig. 2B-C). Hematopoietic stem cells, identified by co-expression of *Kit, Ly6a(Sca1)* and *Flt3* mRNAs (4, 5), clustered together with myeloid and erythroid progenitors (Suppl. Fig. S5A- B). The number of ESC-derived HSCs was higher in BM of mouse-rat chimeras compared to the control (Suppl. Fig. S5C), findings consistent with FACS analysis (Fig. 1H). Only 6 out of 6018 bone marrow cells (0.1%) in mouse-rat chimeras contained both mouse and rat mRNA transcripts (Suppl. Tables S2 and S3), indicating that the fusion of mouse and rat BM cells is rare. Thus, although the cellular composition of ESC-derived hematopoietic BM cells was similar in mouse-rat and mouse-mouse chimeras, mouse-rat bone marrow was enriched in HSCs, LMPPs and CMP-derived erythroid, myeloid and neutrophil progenitors.

### Single cell RNA sequencing identifies close similarities in gene expression signatures between ESC-derived hematopoietic cells in mouse-rat and mouse-mouse chimeras

Comparison of gene expression signatures between mouse-rat and mouse-mouse chimeras revealed significant similarities among ESC-derived hematopoietic cell types. Lymphoid progenitors and pro-B cells isolated from mouse-rat and control chimeras expressed *Ncl, Mif, Rcsd1* and *Tspan13*, whereas pre-B cells expressed *Hmgb2, Tcf3* and *Pgls* (Fig. 3A). *Cd79a* and *CD79b* transcripts were detected in B cells of mouse-rat and control chimeras, whereas *Cd3g* and *Klrd1* were restricted to T cells (Fig. 3A). Based on the correlation analysis, gene expression profiles of all lymphoid cell types were similar between mouse-rat and control chimeras (Fig. 3B). Likewise, gene expression signatures of myeloid, erythroid and neutrophil progenitors and their derivatives in the bone marrow were similar in both experimental groups (Fig. 4A-B). Furthermore, single cell RNAseq identified close similarities in gene expression signatures of ESC-derived HSCs and LMPPs in both chimeras (Suppl. Figs. S5D and S6). Thus, gene expression signatures of ESC-derived hematopoietic cells were similar in mouse-rat and control mouse-mouse chimeras.

**Figure 3.**
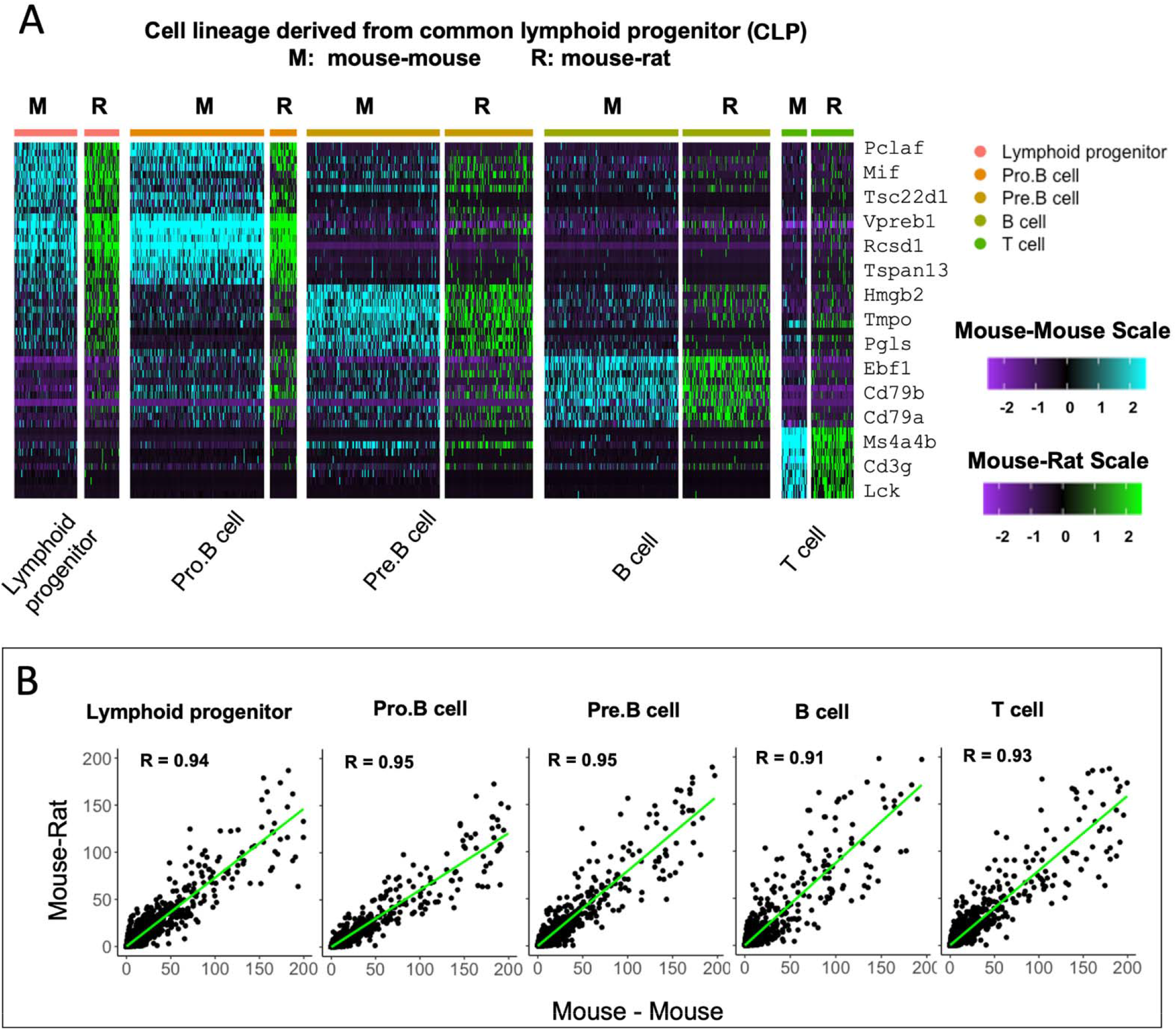
ESC-derived lymphoid cell types in mouse-rat and mouse-mouse chimeras exhibit identical gene expression profiles. ***A,*** Heatmap shows significant similarities in gene expression signatures of lymphoid progenitor cells, Pro-B, Pre-B, B and T cells obtained from mouse-rat (R) and mouse-mouse chimeras (M). Single cell RNAseq was performed using BM cell suspensions that were FACS-sorted for GFP to identify ESC-derived cells. ***B,*** Linear regression analysis shows the correlation index (R) between gene expression profiles in individual lymphoid cell clusters from mouse-rat and mouse-mouse chimeras.

**Figure 4.**
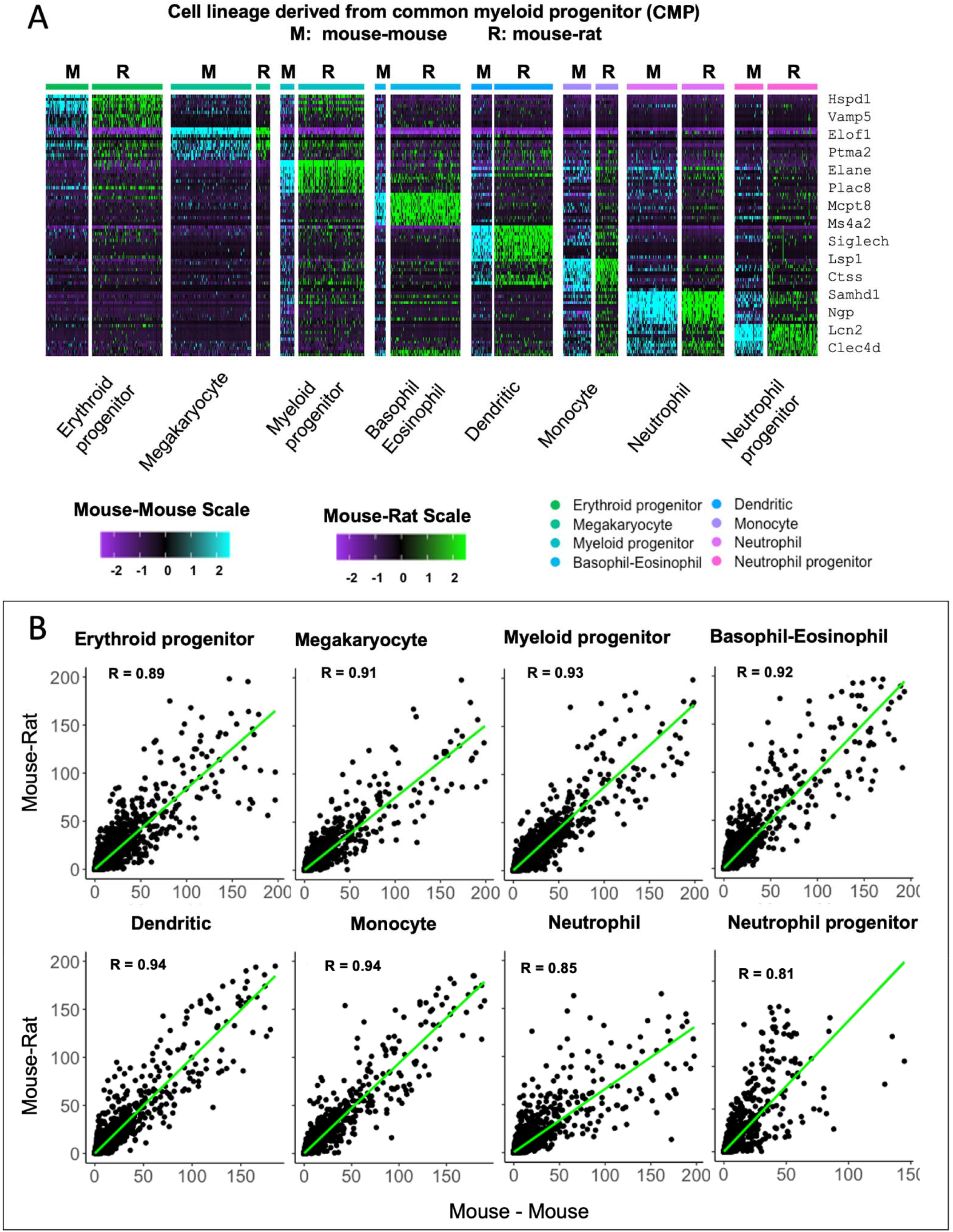
ESC-derived myeloid cell types in mouse-rat and mouse-mouse chimeras exhibit similar gene expression profiles. ***A,*** Heatmap shows significant similarities in gene expression signatures of myeloid progenitor cells, megakaryocytes, erythroid progenitor cells, basophils, eosinophils, neutrophils, dendritic cells, monocytes and neutrophil progenitor cells obtained from mouse-rat (R) and mouse-mouse chimeras (M). Single cell RNAseq was performed using BM cell suspensions that were FACS-sorted for GFP to identify ESC-derived cells. ***B,*** Linear regression analysis shows the correlation index (R) between gene expression profiles in individual myeloid cell clusters from mouse-rat and mouse-mouse chimeras.

### Transplantation of ESC-derived bone marrow cells from interspecies mouse-rat chimeras rescues lethally-irradiated syngeneic mice

To test functional properties of mouse BM hematopoietic progenitor cells derived through a rat, cells were FACS-sorted for GFP from the bone marrow of juvenile mouse-rat chimeras and transferred into the tail vein of syngeneic C57BL/6 adult mice that received the lethal dose of whole-body gamma-irradiation three hours prior to the bone marrow transplant (Fig. 5A). Consistent with published studies (4, 5, 14), all mice without bone marrow transplant died between 9 and 12 days after irradiation (Fig. 5B). In contrast, all 20 mice transplanted with GFP^+^ BM cells from mouse-rat chimeras survived after lethal irradiation (Fig. 5B-C). Blood analysis of mice harvested 8 days after irradiation showed significant decreases in white blood cells (WBC), red blood cells (RBC), platelets (PLT), hemoglobin (Hb) as well as numbers of granulocytes, monocytes and lymphocytes (Fig. 6A-B and Suppl. Figs. S7 and S8). Transplantation of ESC-derived BM cells from mouse-rat chimeras increased WBC and the numbers of granulocytes, monocytes and lymphocytes in the peripheral blood at day 8 (Fig. 6A-B and Suppl. Figs. S7 and S8). Contribution of ESC-derived BM cells to granulocytes, monocytes and B cells was higher compared to erythroid and T cells (Fig. 6C and Suppl. Fig. S9). At 5 months after BM transplantation, ESC-derived cells completely restored blood cell numbers, PLT and Hb in lethally irradiated mice (Fig. 6A-B and Suppl. Figs. S7 and S8). Long-term contributions of ESC-derived BM cells to all hematopoietic cell lineages in the peripheral blood were between 49% and 96% (Fig. 6D and Suppl. Fig. S9). Thus, transplantation of ESC-derived bone marrow cells from mouse-rat chimeras prevented mortality and restored hematopoietic blood lineages in lethally irradiated syngeneic mice.

**Figure 5.**
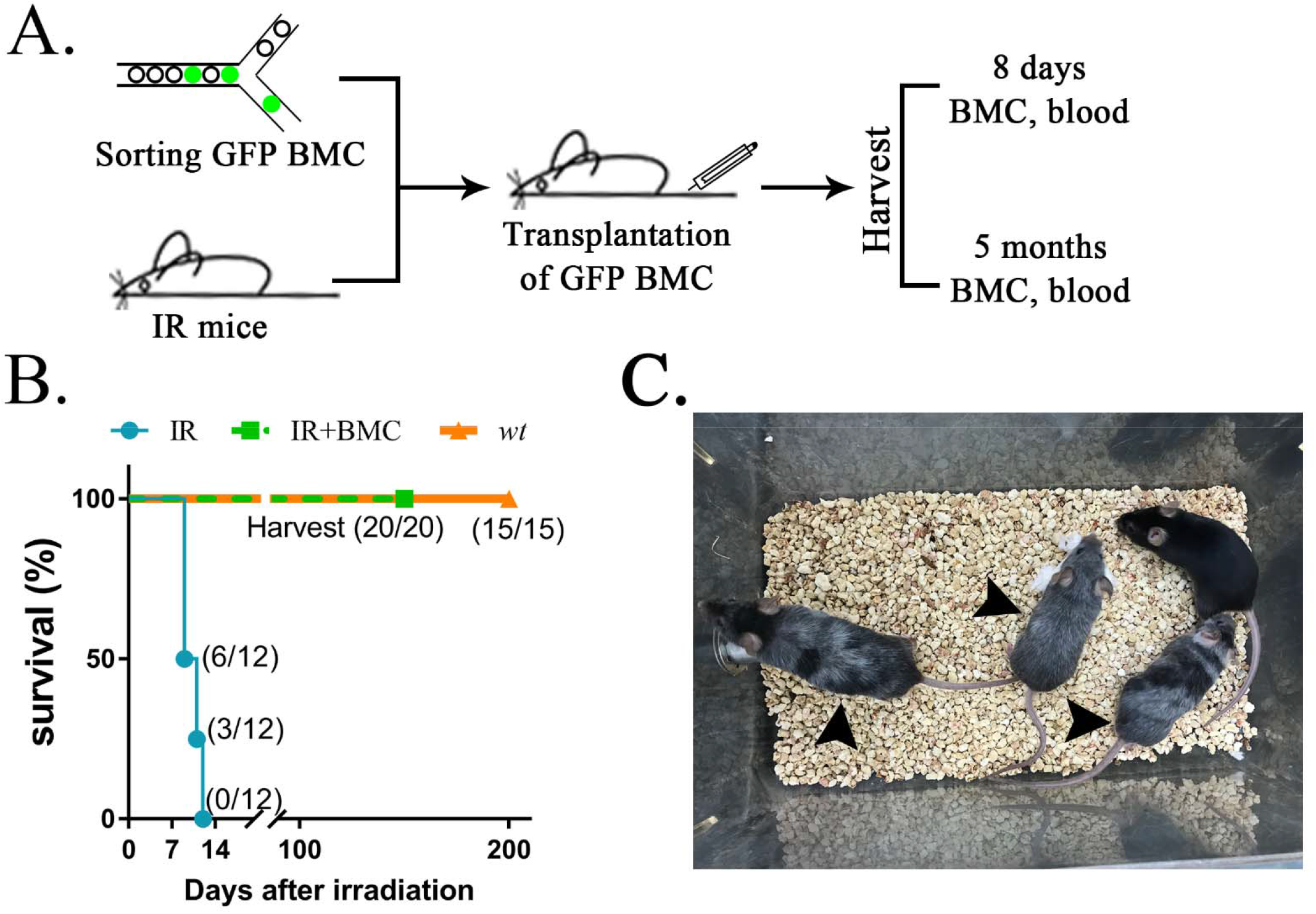
Transplantation of mouse ESC-derived BM cells from interspecies mouse-rat chimeras rescues lethally irradiated syngeneic mice. ***A,*** Schematic diagram shows transplantation of ESC-derived bone marrow cells (BMC) into lethally irradiated (IR) mice. ESC- derived cells were obtained from the bone marrow of juvenile mouse-rat chimeras using FACS-sorting for GFP^+^ cells. Bone marrow and peripheral blood were harvested 8 days and 5 months after BM transplantation. *B,* Kaplan-Meier survival analysis shows a 100% mortality in irradiated mice. Survival is dramatically improved after transplantation of irradiated mice with ESC-derived BM cells obtained from mouse-rat chimeras (IR + BMC). Survival in untreated wild type (wt) mice is shown as a control (n=12-20 mice in each group). *C,* Photograph shows irradiated C57BL/6 mice 5 months after successful bone marrow transplantation. Untreated C57BL/6 mouse is shown as a control. Grey color of irradiated mice (arrows) is consistent with large doses of whole-body radiation treatment.

**Figure 6.**
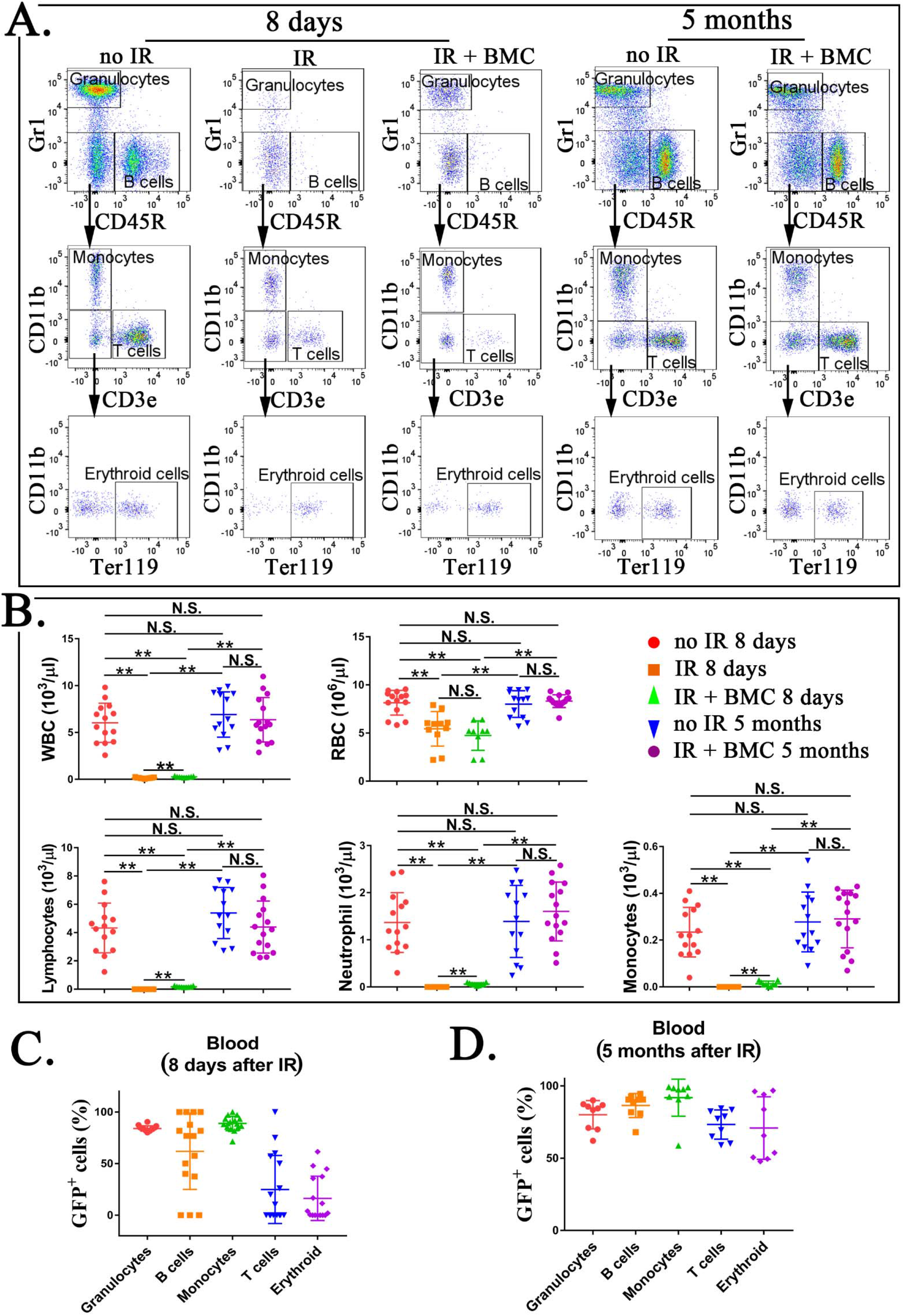
Transplantation of mouse ESC-derived BM cells from interspecies mouse-rat chimeras restores blood hematopoietic cell lineages in lethally irradiated syngeneic mice. ***A,*** FACS analysis shows identification of granulocytes, B cells, monocytes, T cells and erythroid cells in the peripheral blood 8 days and 5 months after BM transplantation. Blood samples were obtained from untreated mice (no IR), lethally irradiated mice without bone marrow transplant (IR), and lethally irradiated mice with bone marrow transplant (IR+BMC). BM transplantation was performed using ESC-derived BM cells obtained from juvenile mouse-rat chimeras. ***B,*** Blood analysis shows that transplantation with ESC-derived BM cells from mouse-rat chimeras increases white blood cell (WBC) counts and red blood cell (RBC) counts in the peripheral blood. Concentrations of lymphocytes, monocytes and neutrophil in the blood were also increased after BM transplantation (n=9-15 mice in each group). ***C-D,*** FACS analysis for GFP^+^ cells in each cell subset shows that ESC-derived BM cells from mouse-rat chimeras contribute to multiple hematopoietic cell lineages in the peripheral blood of lethally irradiated mice (n=9-16 mice in each group), p<0.01 is **, N.S. indicates no significance.

### Transplantation of ESC-derived bone marrow cells from interspecies mouse-rat chimeras resulted in the long-term contribution of donor cells to hematopoietic progenitor cells

Based on FACS analysis of irradiated mice at day 8, whole-body irradiation decreased the number of hematopoietic progenitor cells in the bone marrow, including LSKs, ST-HSCs and LT-HSCs (Fig. 7A-B and Suppl. Fig. S10). Transplantation of ESC-derived BM cells significantly increased LSKs but did not affect the numbers of ST-HSCs and LT-HSCs in irradiated mice (Fig. 7A-B and Suppl. Fig. S10). Contribution of ESC-derived BM cells to Lin^−^ and LSK cell subsets was high, whereas ESC contribution to ST-HSCs and LT-HSCs at day 8 was low (Fig. 7D and Suppl. Fig. S11). At 5 months after BM transplantation, percentages of LSKs, ST-HSCs and LT-HSCs in the bone marrow were significantly increased (Fig. 7A-B and Suppl. Fig. S10). Long-term contribution of ESC-derived BM cells to LSKs, ST-HSCs and LT-HSCs was between 92% and 95% (Fig. 7E and Suppl. Fig. S11). Altogether, transplantation of ESC-derived bone marrow cells from mouse-rat chimeras resulted in efficient, long-term contribution of donor cells to the bone marrow and blood of lethally irradiated mice.

**Figure 7.**
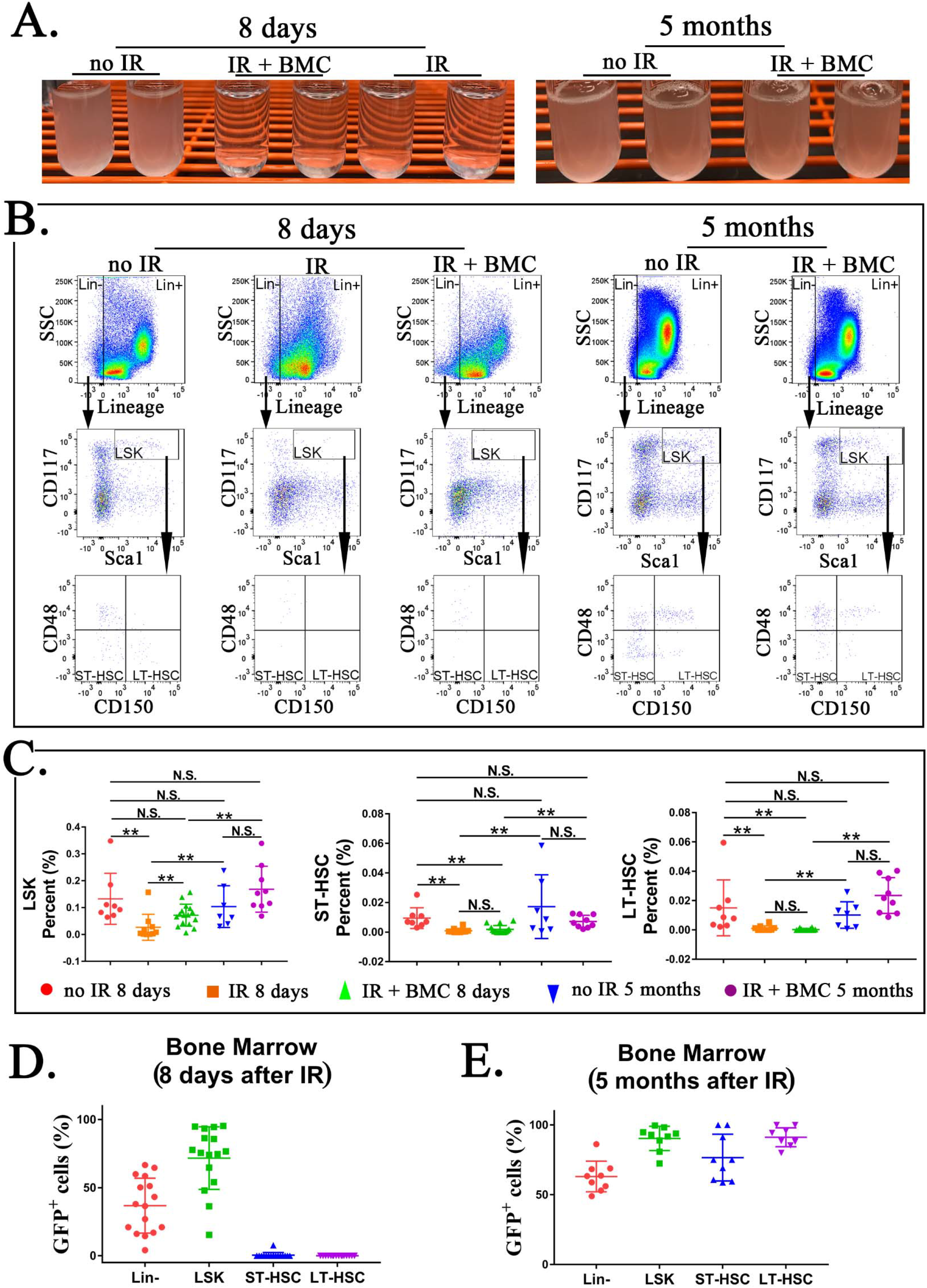
Transplantation of mouse ESC-derived BM cells from interspecies mouse-rat chimeras restores hematopoietic progenitor cells in the bone marrow of irradiated syngeneic mice. ***A***, Photographs show cell suspensions obtained from the bone marrow of untreated mice (no IR), lethally irradiated mice without bone marrow transplant (IR), and lethally irradiated mice with bone marrow transplant (IR+BMC). Mice were harvested 8 days (left image) or 5 months after BM transplantation (right image). ESC-derived BM cells from juvenile mouse- rat chimeras were used for BM transplantation. ***B***, FACS analysis shows identification of Lineage^−^ (Lin^−^) cells, LSKs, ST-HSCs and LT-HSCs in the bone marrow of irradiated mice 8 days and 5 months after BM transplantation. ***C***, FASC analysis shows that transplantation with ESC-derived BM cells from mouse-rat chimeras increases percentages of LSKs, ST-HSCs and LT-HSCs in the bone marrow of irradiated mice 5 months after BM transplantation (n=9-16 mice in each group). ***D-E***, ESC-derived BM cells from mouse-rat chimeras contribute to multiple hematopoietic progenitor cells in the bone marrow of irradiated mice (n=9-16 mice in each group), p<0.01 is **, N.S. indicates no significance.

## DISCUSSION

Recent single cell RNA sequencing studies identified remarkable diversity of hematopoietic cell types in the bone marrow (1). Generation of functional bone marrow cells from pluripotent ESCs or iPSCs in a dish or in organoids represents a formidable challenge (4, 5). In the present study, we used blastocyst complementation to generate a diversity of hematopoietic cell types from mouse ESCs in rat embryos. Interspecies mouse-rat chimeras were viable and contained approximately 25% of ESC-derived mouse cells in the bone marrow. It is possible that inactivation of genes critical for hematopoiesis in rat embryos prior to blastocyst complementation can improve the integration of mouse ESCs into the bone marrow of mouse-rat chimeras. This approach was supported by recent studies with mouse-mouse chimeras, in which ESCs contributed to more than 90% of hematopoietic cells in mice deficient for either *Kit* or *Flk1* (28, 34). While ESCs contributed to all hematopoietic cell lineages in interspecies bone marrow, the percentage of lymphoid progenitors was lower, whereas the percentages of myeloid progenitor cells and HSCs were higher in mouse-rat chimeras compared to control mouse-mouse chimeras. Since both chimeras were produced by complementing blastocysts with mouse ESCs from the same ESC-GFP cell line, it is unlikely that these changes are dependent on donor ESCs. It is possible that the observed differences in BM cellular composition between mouse-rat and mouse-mouse chimeras are due to interactions of donor ESCs with the host embryo. Structural and functional differences between hormones, growth factors and their receptors in rats and mice can contribute to the efficiency or timing of differentiation of mouse ESCs into hematopoietic cell lineages in BM of chimeras.

Despite mosaicism in interspecies bone marrow, mouse ESC-derived cells from multiple hematopoietic cell lineages were highly differentiated and indistinguishable from the normal mouse bone marrow cells based on gene expression signatures and cell surface proteins. Consistent with functional competency of ESC-derived bone marrow cells, transplantation of these cells into lethally-irradiated syngeneic mice prevented mortality and resulted in long-term contribution of ESC-derived cells to all hematopoietic cell lineages in the bone marrow and peripheral blood. Our results are consistent with recent studies demonstrating the ability of mouse ESCs to generate functional pancreatic, endothelial and kidney cells in interspecies mouse-rat chimeras (25, 27, 35). Interestingly, long-term contribution of donor BM cells to ST- HSCs and LT-HSCs of irradiated mice was high, supporting the ability of donor HSCs to self- renew. In contrast, the short-term contribution of donor BM cells to ST-HSCs and LT-HSCs of irradiated mice was low. Low contribution of donor BM to HSCs at day 8 is not surprising considering an acute hematopoietic deficiency in lethally irradiated mice. It is possible that a majority of donor-derived HSCs undergo rapid differentiation into other hematopoietic cell types to compensate for the loss of injured hematopoietic cells after irradiation.

Generation of intraspecies chimeras through blastocyst complementation creates an interesting opportunity to use patient-derived iPSCs to produce tissues or even organs in large animals, for example, pigs or sheep, which can serve as “biological reactors”. However, at this stage of technological advances it is impossible to restrict the integration of ESC/iPSC-derived cells into selected organs or cell types. Off-target integration of ESCs and iPSCs into the brain, testes and sensory organs raises important ethical concerns for the use of human-animal chimeras in regenerative medicine (53, 54). To improve the selectivity of ESC/iPSC integration into chimeric tissues, various genetic modifications can be introduced into the host embryos to advance the technology. Harvest of tissues from chimeric embryos instead of adult chimeras can alleviate some of the ethical concerns, suggesting a possibility of using chimeric embryos as a potential source of patient-specific hematopoietic progenitor cells.

In summary, blastocyst complementation of rat embryos with mouse ESCs was used to simultaneously generate all hematopoietic cell lineages in the bone marrow. ESC-derived cells in mouse-rat chimeras were indistinguishable from normal mouse hematopoietic BM cells based on gene expression signatures and cell surface markers. Transplantation of ESC-derived BM cells rescued lethally-irradiated syngeneic mice and resulted in long-term contribution of donor cells to hematopoietic cell lineages. Thus, the interspecies chimeras could be considered for *in vivo* differentiation of patient-derived iPSCs into hematopoietic cell lineages for future cell therapies.

## ACKNOWLEDGMENTS

We thank Mrs. Erika Smith for excellent secretarial support. This work was supported by NIH Grants HL141174 (to V.V.K.), HL149631 (to V.V.K.) and HL152973 (to V.V.K and T.V.K.).

## AUTHOR CONTRIBUTIONS

B.W. and V.V.K., designed the study; B.W., G.W. and E.L., conducted experiments; G.W., conducted bioinformatic analyses; B.W., G.W., E.L., O.A.K., Z.T., S.D., and T.V.K., analyzed the data and provided critical insights; B.W., O.A.K. and V.V.K., wrote the manuscript with input from all authors.

---

**Supplemental Figure S1.**
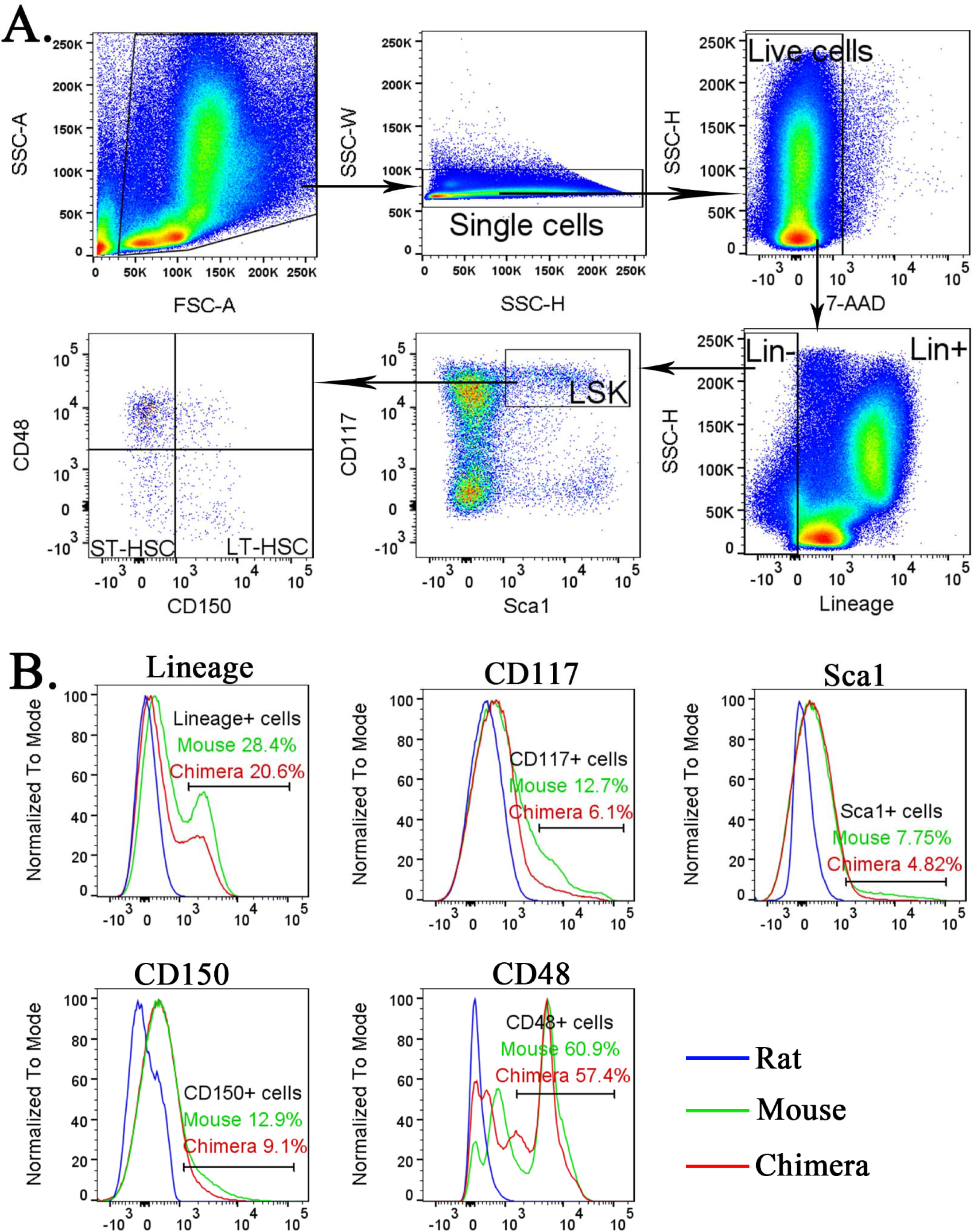
Identification of lineage^−^ cells, LSKs, ST-HSCs and LT-HSCs in the bone marrow. ***A,*** FACS gating strategy shows the identification of lineage^−^ (Lin^−^) cells, LSKs, ST-HSCs and LT-HSCs in the bone marrow of *wt* mice. ***B,*** Histograms show specificity of antibodies against *Lineage* antigens, CD117 (c-KIT), Sca1, CD150 and CD48. To identify mouse Lin^−^ cells, LSKs, ST-HSCs and LT-HSCs, cell suspensions from BM of *wt* mouse, mouse-rat chimera and *wt* rat were compared to determine the gating.

**Supplemental Figure S2.**
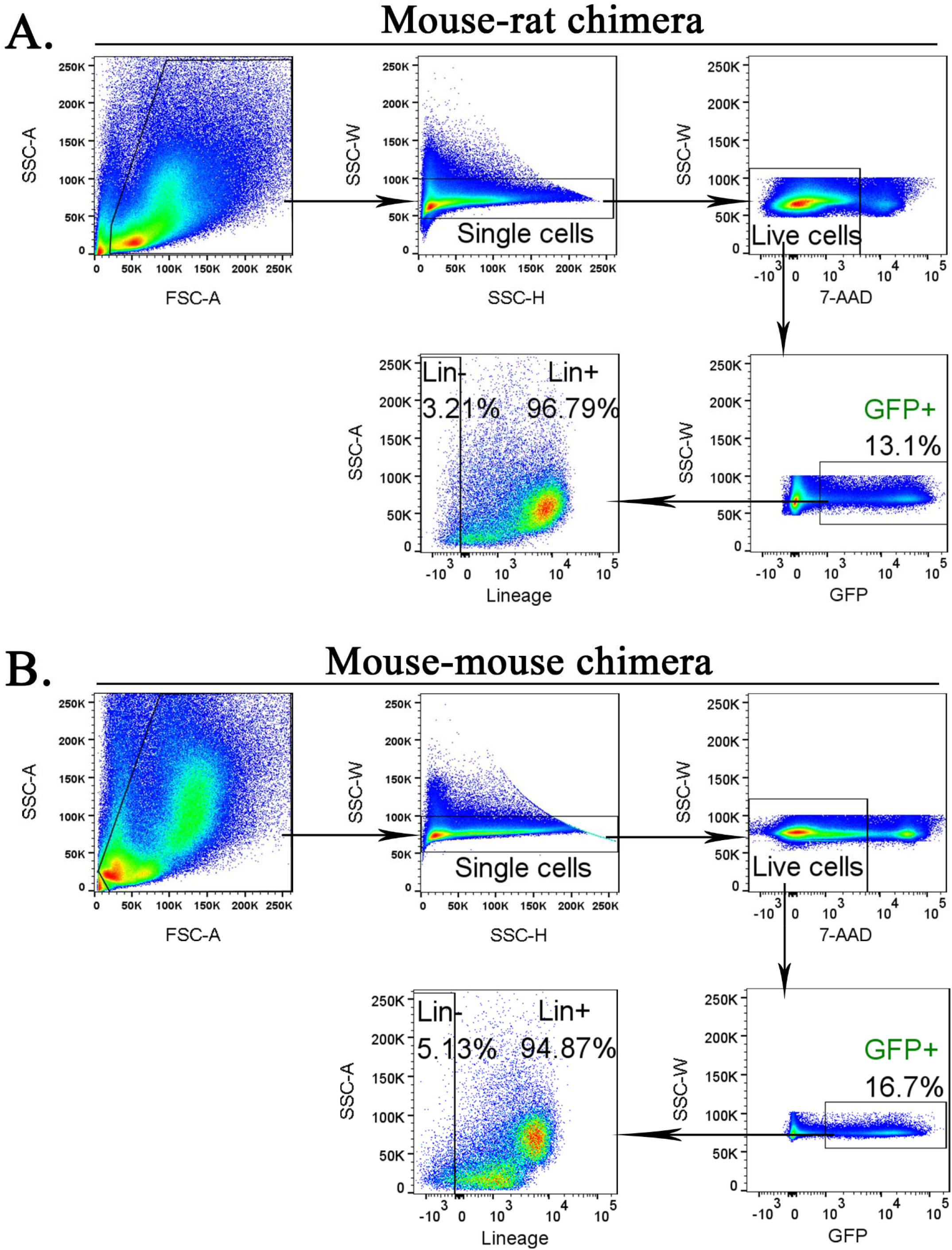
Purification of mouse ESC-derived cells from bone marrow of mouse-rat and mouse-mouse chimeras before scRNAseq. FACS gating strategy shows identification of ESC-derived Lin^−^ and Lin^+^ cell subsets in the bone marrow of mouse-rat (**A**) and mouse-mouse chimeras (**B**). Chimeric BM cells were harvested at P10.

**Supplemental Figure S3.**
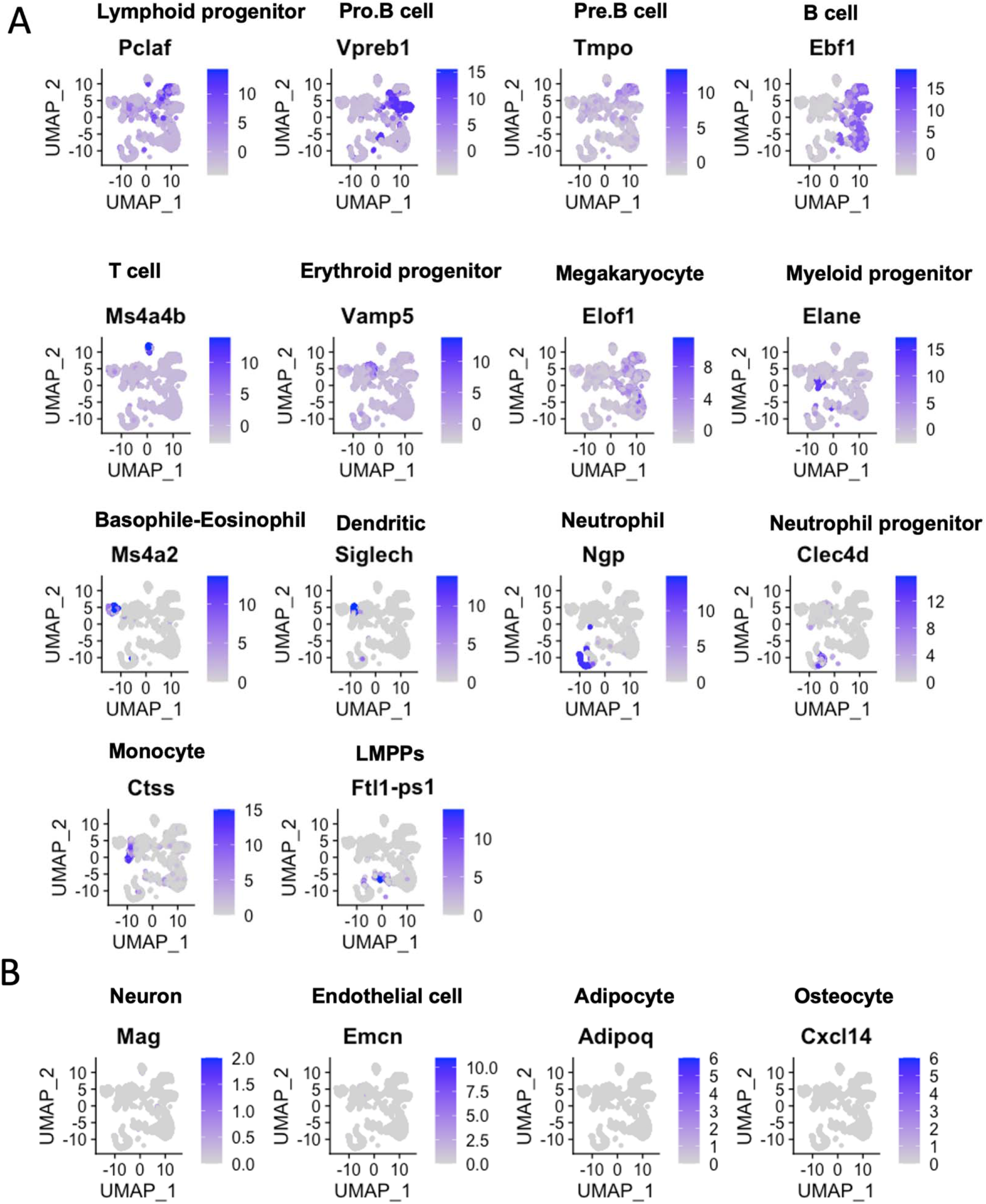
Single cell RNAseq analysis identifies hematopoietic cell subsets in the bone marrow of mouse-rat chimeras. ***A,*** The integrated projection of ESC-derived BM hematopoietic cells from mouse-rat and mouse-mouse (control) chimeras. Cells were obtained from the bone marrow of P10 chimeras. Cell clusters were identified using Uniform Manifold Approximation and Projection (UMAP) method. Expression of marker genes shows different hematopoietic cell clusters in the bone marrow. ***B,*** Genes enriched in neurons, endothelial cells, adipocytes and osteocytes are not detected in cell clusters from chimeric bone marrow.

**Supplemental Figure S4.**
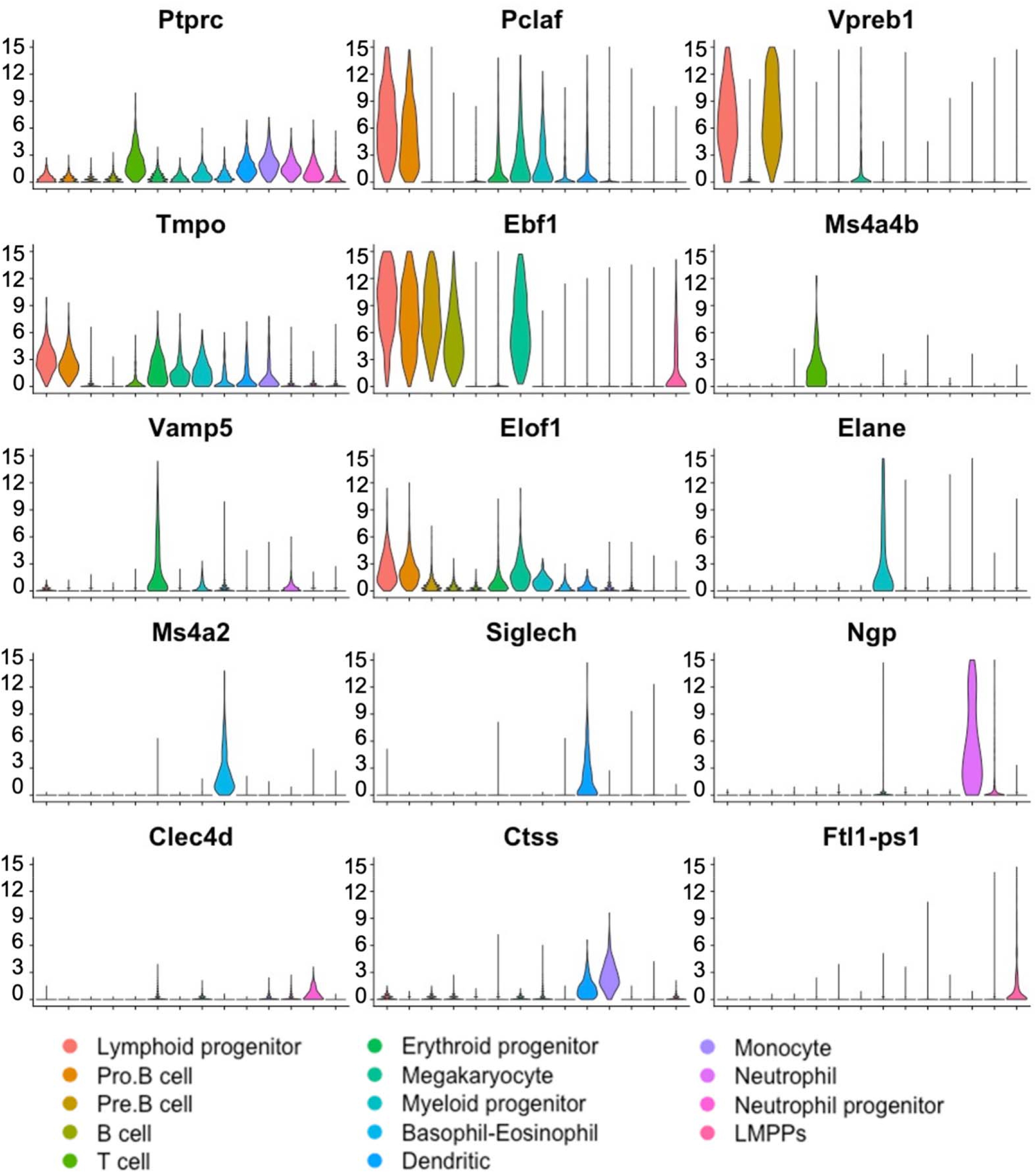
Violin plots confirm expression of hematopoietic marker genes in BM cell clusters. Single cell RNAseq was performed using ESC-derived BM hematopoietic cells from mouse-rat and mouse-mouse P10 chimeras. Cell clusters were identified using UMAP. Violin plots show expression of selected hematopoietic genes in BM cell clusters. *Ptprc (Cd45)* mRNA is expressed in all hematopoietic cell types.

**Supplemental Figure S5.**
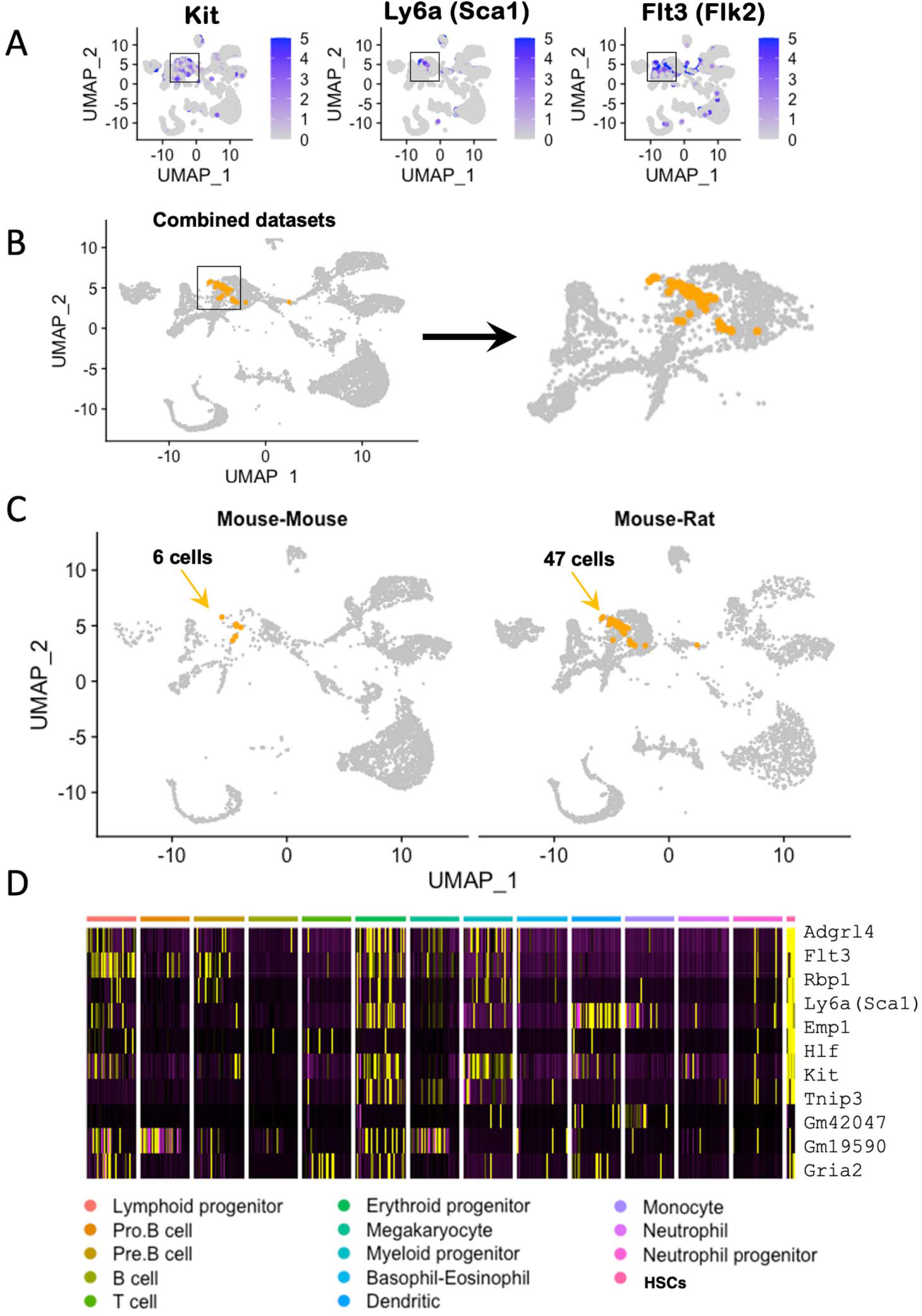
Single cell RNAseq analysis identifies genes expressed in hematopoietic stem cells in chimeric bone marrow. ***A,*** UMAP analysis shows expression of genes enriched in hematopoietic stem cells (HSCs), including *Kit*, *Ly6a (Sca-1)* and *Flt3 (Flk2)*, using the integrated projection of ESC-derived BM hematopoietic cells from mouse-rat and mouse-mouse (control) chimeras. Cells were obtained from the bone marrow of P10 chimeras. ***B,*** *Kit^+^Ly6a^+^Flt3^+^* triple positive cells are identified in myeloid and erythroid progenitor cell clusters in the combined scRNAseq dataset. ***C,*** Separate views of triple positive cells in individual scRNAseq datasets show increased number of HSCs in mouse-rat chimera compared to mouse-mouse chimera. ***D,*** Heatmap shows expression of genes enriched in HSCs.

**Supplemental Figure S6.**
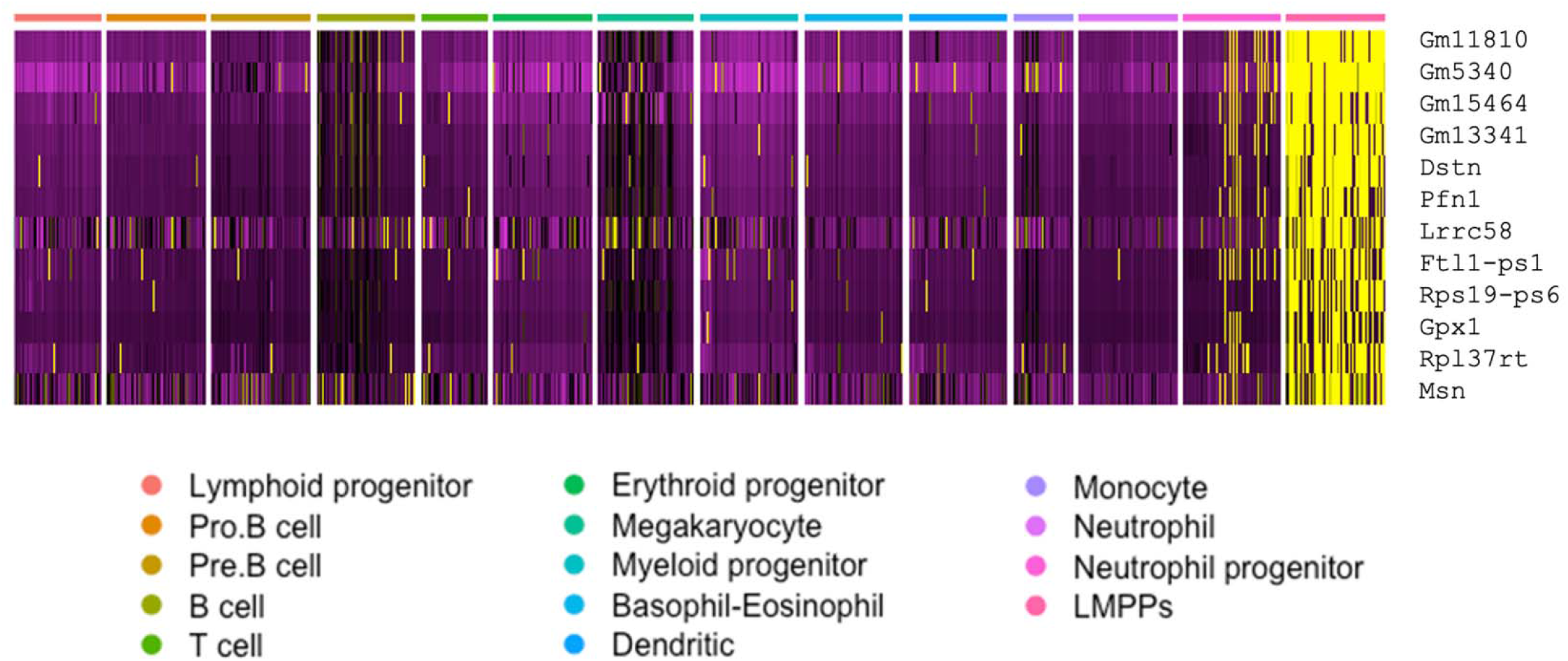
Heatmap identifies gene expression profile of ESC-derived lymphoid-primed multipotent progenitors from mouse-rat and mouse-mouse chimeras. Combined analysis of ESC-derived BM hematopoietic cells from mouse-rat and mouse-mouse chimeras compares gene expression signature of lymphoid-primed multipotent progenitor cells (LMPPs) with gene expression signatures of other myeloid and lymphoid BM cells. Single cell RNAseq was performed using BM cell suspensions from P10 chimeras. ESC-derived cells were purified using FACS.

**Supplemental Figure S7.**
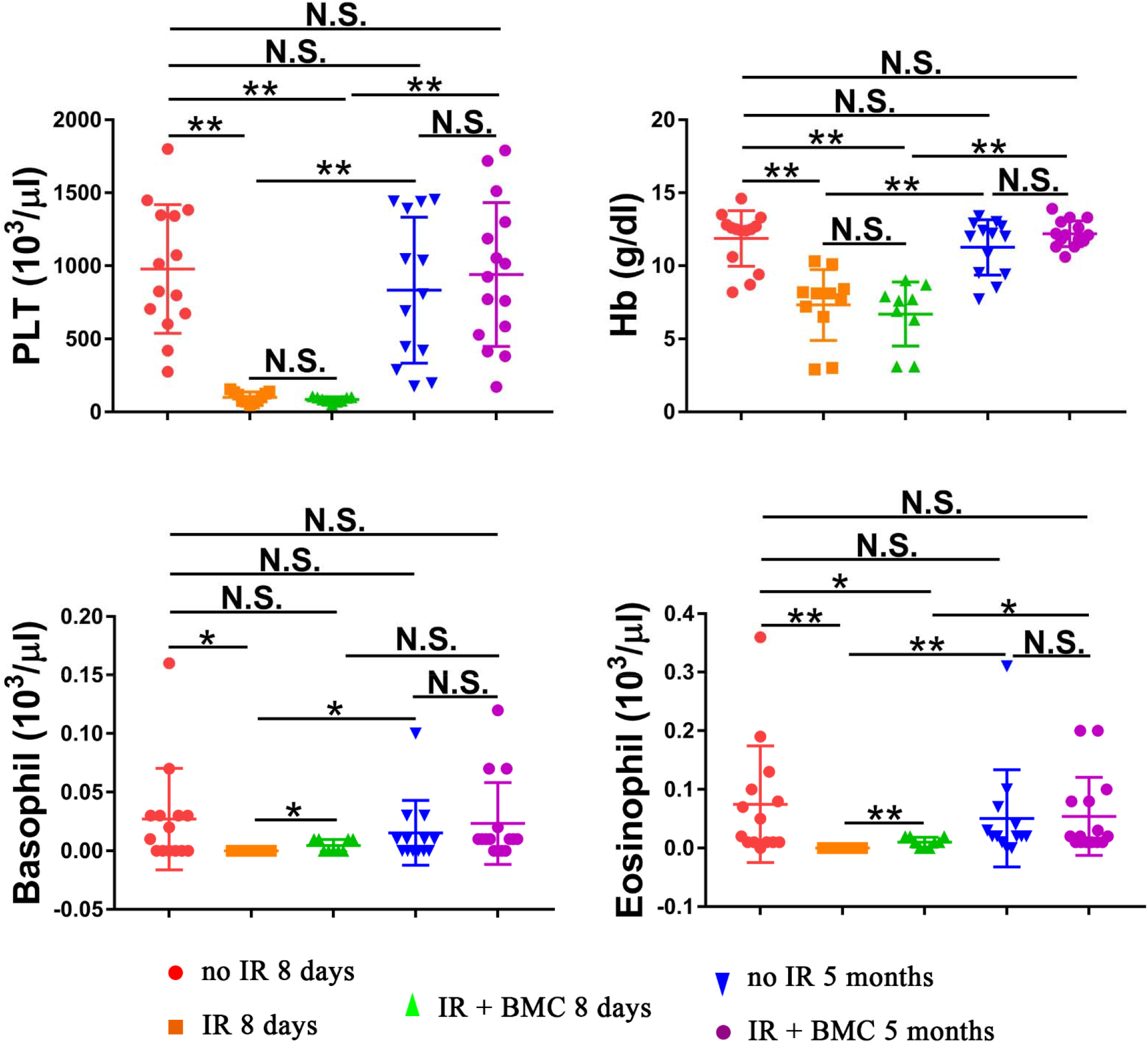
Transplantation of irradiated mice with ESC-derived BM cells from mouse-rat chimeras increases concentrations of platelets, hemoglobin, basophils and eosinophils in the peripheral blood. Blood samples were obtained from untreated mice (no IR), lethally irradiated mice without bone marrow transplant (IR), and lethally irradiated mice with bone marrow transplant (IR+BMC). BM transplantation was performed using ESC-derived BM cells obtained from juvenile mouse-rat chimeras. Mice were harvested 8 days or 5 months after BM transplantation. Concentrations of basophils and eosinophils in the peripheral blood were significantly increased 8 days after BM transplantation. Concentrations of platelets (PLT), hemoglobin (Hb), basophils and eosinophils were fully restored 5 months after BM transplantation (n=9-15 mice in each group), p<0.05 is *, p<0.01 is **, N.S. indicates no significance.

**Supplemental Figure S8.**
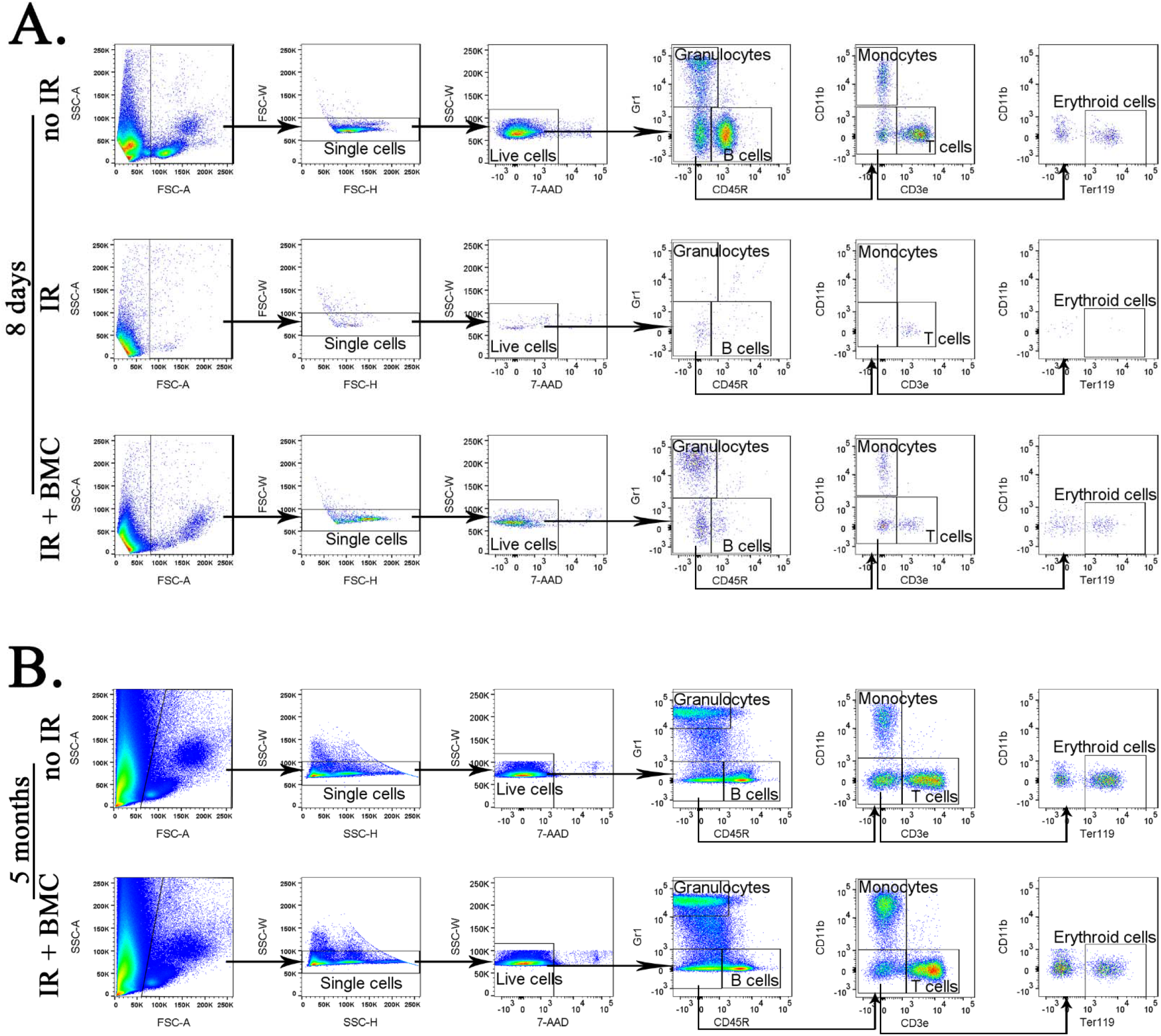
FACS analysis shows identification of granulocytes, B cells, monocytes, T cells and erythroid cells in the peripheral blood of irradiated mice after BM transplantation. BM transplantation was performed using ESC-derived BM cells that were purified from juvenile mouse-rat chimeras. Blood samples were obtained from untreated mice (no IR), lethally irradiated mice without bone marrow transplant (IR), and lethally irradiated mice with bone marrow transplant (IR+BMC). Blood was harvested 8 days (**A**) or 5 months after BM transplantation (**B**) and used for FACS analysis.

**Supplemental Figure S9.**
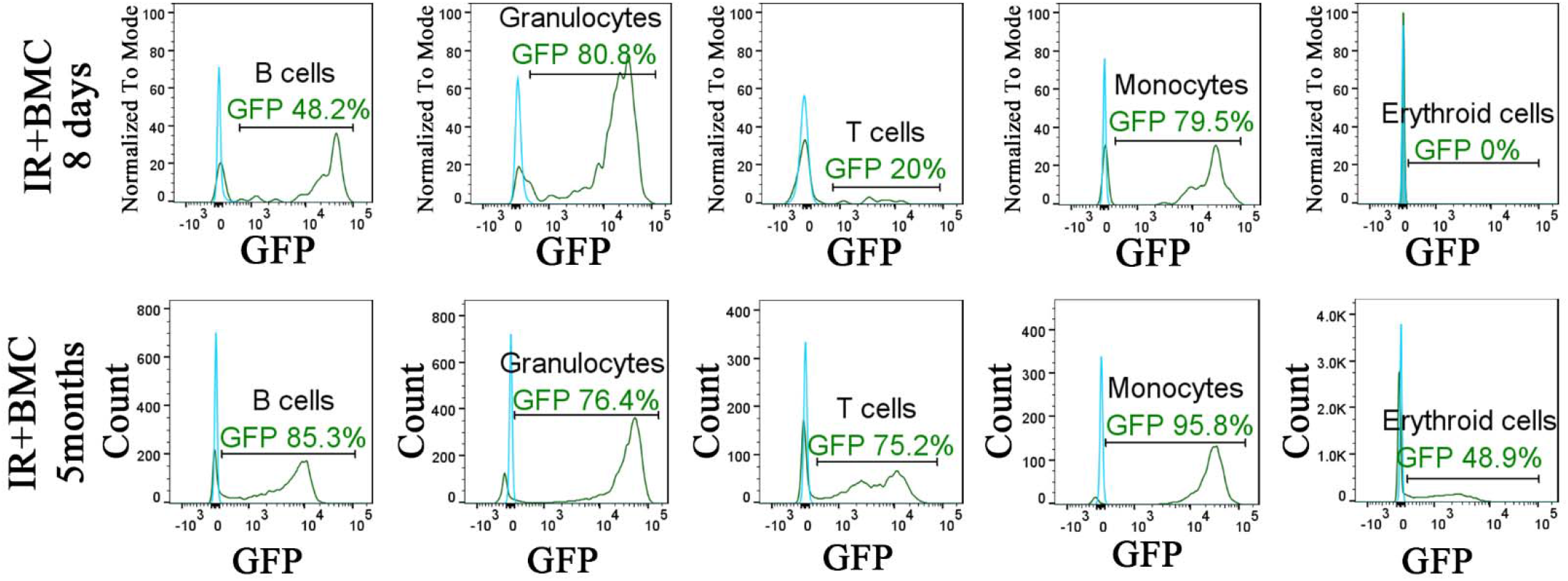
Identification of ESC-derived cells in the peripheral blood of irradiated mice after BM transplantation. Histograms show the presence of ESC-derived (GFP- positive) granulocytes, B cells, T cells, monocytes and erythroid cells in the peripheral blood of irradiated mice after BM transplantation (green line). Blood of mice without BM transplantation is used to identify autofluorescence in GFP channel (blue line). Blood samples were harvested 8 days or 5 months after BM transplantation and used for FACS analysis.

**Supplemental Figure S10.**
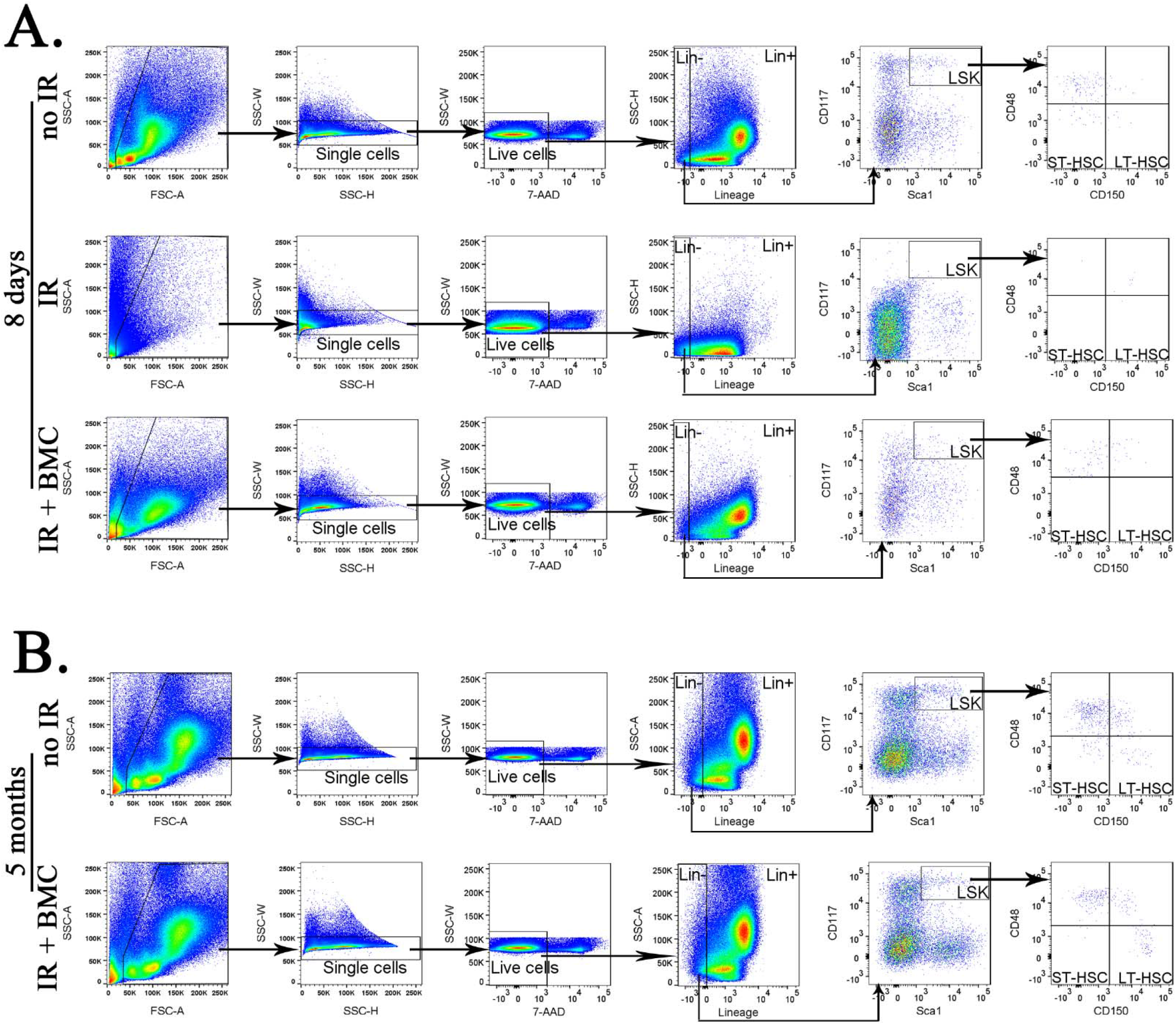
FACS analysis shows identification of Lin^−^ cells, LSKs, ST-HSCs and LT-HSCs in the bone marrow of irradiated mice 8 days and 5 months after BM transplantation. BM transplantation was performed using ESC-derived BM cells that were purified from juvenile mouse-rat chimeras. Bone marrow was obtained from one tibia and one fibula bones of untreated mice (no IR), lethally irradiated mice without bone marrow transplant (IR), and lethally irradiated mice with bone marrow transplant (IR+BMC). Bone marrow was harvested 8 days (**A**) or 5 months after BM transplantation (**B**) and used for FACS analysis.

**Supplemental Figure S11.**
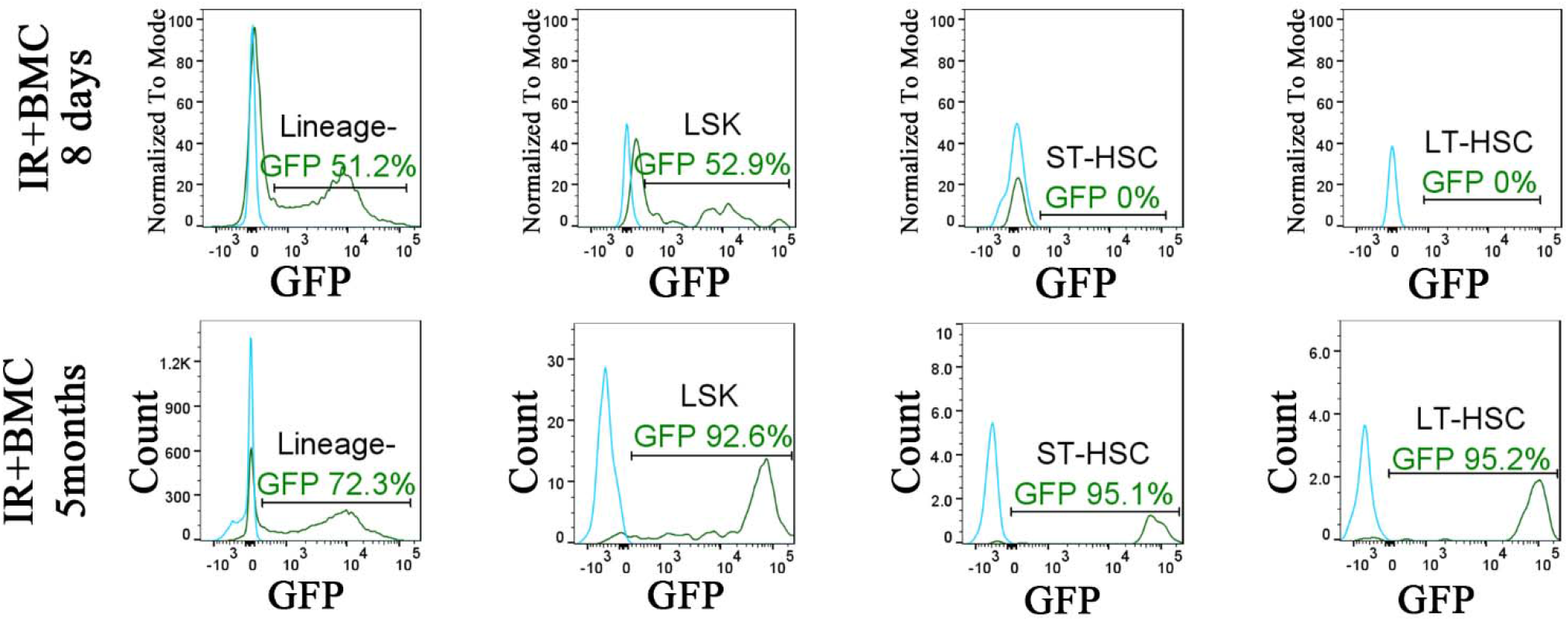
Identification of ESC-derived hematopoietic cells in the bone marrow of irradiated mice after BM transplantation. Histograms show the presence of ESC-derived (GFP-positive) Lin^−^ cells, LSKs, ST-HSCs and LT-HSCs in the bone marrow of irradiated mice after BM transplantation (green line). For each cell subset, bone marrow of mice without BM transplantation is used to identify autofluorescence in GFP channel (blue line). Bone marrow was obtained from one tibia and one fibula bones 8 days or 5 months after BM transplantation.

**Supplemental Table S1.** Antibodies used for FACS analysis.

| Antibody | Manufacturer | Catalog No. | Dilution |
| --- | --- | --- | --- |
| Lineage- | BD Bioscience | 561317 | 1:100 |
| CD117 | Thermo Fisher | 12-1171-83 | 1:100 |
| Sca1 | Thermo Fisher | 17-5981-81 | 1:100 |
| CD150 | Thermo Fisher | 25-1502-80 | 1:100 |
| CD45R | Thermo Fisher | 83-0452-41 | 1:100 |
| Gr1 | Biolegend | 108433 | 1:100 |
| CD3e | Biolegend | 100312 | 1:100 |
| CD11b | Biolegend | 101216 | 1:100 |
| Ter119 | Biolegend | 116228 | 1:100 |
| CD48 | Thermo Fisher | 48-0481-82 | 1:100 |

**Supplemental Table S2.** The number and percentage of hematopoietic BM cells containing both mouse and rat mRNAs (hybrid cells).

| Cluster | Cell number | Cells with mouse and rat mRNAs | Frequency (%) |
| --- | --- | --- | --- |
| Lymphoid progenitor | 199 | 0 | 0 |
| Pro.B cell | 156 | 0 | 0 |
| Pre.B cell | 541 | 0 | 0 |
| B cell | 808 | 4 | 0.0049505 |
| T cell | 243 | 0 | 0 |
| Erythroid progenitor | 1122 | 1 | 0.00089127 |
| Megakaryocyte | 101 | 0 | 0 |
| Myeloid progenitor | 465 | 0 | 0 |
| Basophil-Eosinophil | 543 | 0 | 0 |
| Dendritic | 419 | 0 | 0 |
| Monocyte | 165 | 0 | 0 |
| Neutrophil | 300 | 1 | 0.00333333 |
| Neutrophil progenitor | 358 | 0 | 0 |
| Lineage-negative | 598 | 0 | 0 |

**Supplemental Table S3.**
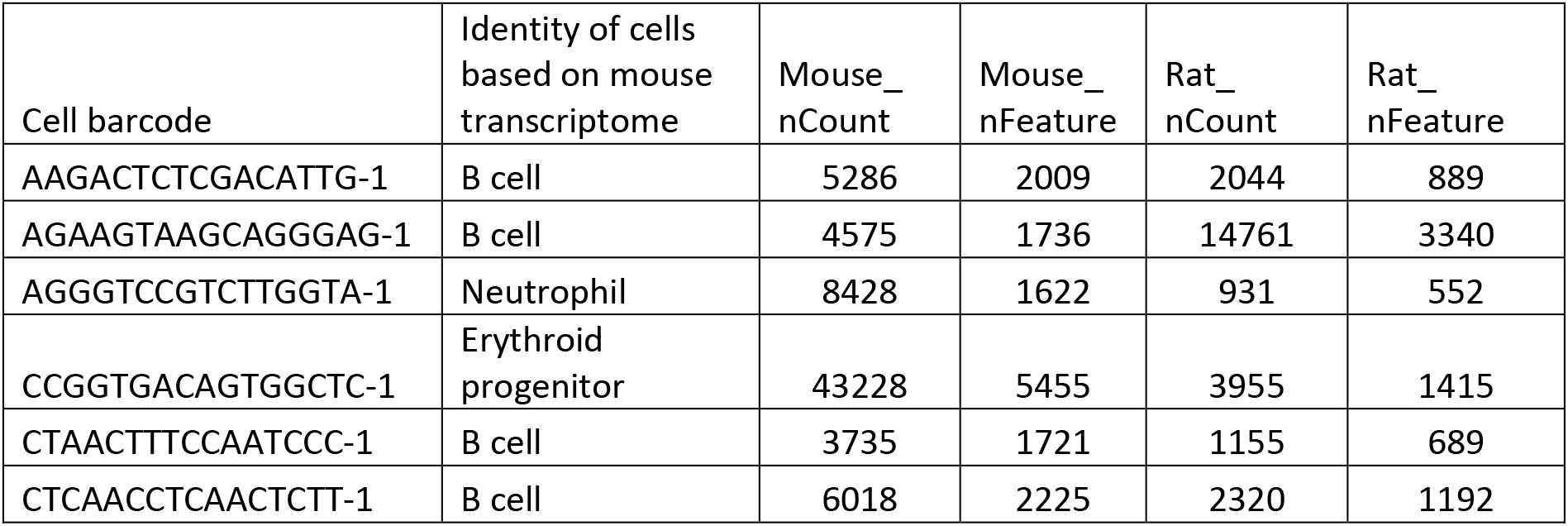
The number of counts and features (genes) in 6 hybrid cells identified in mouse-rat chimera.

